# Anti-Malaria Antibody Engineering Broadens Recognition Motifs and Reveals New Homotypic Interactions that Enhance Protective Breadth

**DOI:** 10.1101/2025.09.30.679367

**Authors:** Jihwan Chun, Prabhanshu Tripathi, Yevel Flores-Garcia, Bharat Madan, Gene A. Lee, Ahmed S. Fahad, Haotian Lei, I-Ting Teng, Nicholas K. Hurlburt, Barbara J. Flynn, Marie Pancera, Kazutoyo Miura, Tongqing Zhou, Azza H. Idris, Fidel Zavala, Robert A. Seder, Peter D. Kwong, Brandon J. DeKosky

**Affiliations:** The Ragon Institute of Massachusetts General Hospital, Massachusetts Institute of Technology, and Harvard University, Cambridge, MA, USA; Department of Chemical Engineering, Massachusetts Institute of Technology, Cambridge, MA, USA; Vaccine Research Center, National Institute of Allergy and Infectious Diseases, National Institutes of Health, Bethesda, MD, USA; Malaria Research Institute, Bloomberg School of Public Health, Johns Hopkins University, Baltimore, MD, USA; Research Technologies Branch, National Institute of Allergy and Infectious Diseases, National Institutes of Health, Bethesda, MD 20892, USA; Vaccine and Infectious Disease Division, Fred Hutchinson Cancer Center, Seattle, WA, USA; Department of Biochemistry and Molecular Biophysics, Columbia University Vagelos College of Physicians and Surgeons, New York, NY, USA; Aaron Diamond AIDS Research Center, Columbia University Vagelos College of Physicians and Surgeons, New York, NY, USA

**Keywords:** malaria, circumsporozoite protein, antibody, homotypic interactions

## Abstract

The monoclonal antibody L9 mediates high-level protection against malaria in children for up to 6 months in Africa. L9 preferentially binds with high affinity to the NVDP minor repeat on the *P. falciparum* circumsporozoite protein (PfCSP). Here, we sought to improve the affinity of L9 to enhance protection against rare strains with two spatially separated minor repeats or a single minor repeat. Site saturation mutagenesis and yeast display-screening identified a panel of affinity-improved variants. *In vivo* challenge showed one variant, L9_yd19, to be modestly more potent against a chimeric transgenic *Plasmodium* encoding PfCSP with two widely spaced minor repeats from a Kenyan parasite strain, with no loss in potency against the benchmark 3D7 strain with its standard complement of minor repeats. L9_yd19 also had high affinity against NANP major repeats and was protective against transgenic *Plasmodium* with PfCSP containing only NANP major repeats (NANP_12_). Cryo-EM studies revealed L9_yd19 to recognize PfCSP with two distinct homotypic interfaces, which combined to yield two trimeric layers of antibodies comprising asymmetric trimers that dimerized in a head-to-head fashion. These data reveal a new antibody mechanism that utilizes interfaces involving dual homotypic symmetry elements, a 2-fold and an asymmetric 3-fold, for potentially improved malaria prevention.

**HIGHLIGHTS:**

- L9 is a highly protective antimalarial antibody that preferentially binds the NVDP minor repeat on *Plasmodium falciparum* circumsporozoite protein (PfCSP) and also binds with low affinity to the NANP major repeat; due to these targeting preferences, it has shown reduced protection against designed transgenic malaria strains with only a single NVDP motif (Fig. 1).
- Using yeast display, a panel of L9 variants were generated based on higher affinity against the minor NVDP and major NANP motifs to determine if they could improve protection against strains with fewer minor repeat regions or only containing major repeats (Figs 1-4).
- One L9 variant, L9_yd19 showed enhanced protection against chimeric transgenic CSP variants with a single minor repeat or two minor repeats in which the spacing was separated; L9_yd19 also showed protection against chimeric transgenic CSP variants containing only the NANP major repeat (Figs. 4-5).
- Cryo-EM analyses revealed L9_yd19 recognition of CSP to comprise two distinct homotypic interfaces: a side-to-side interface within asymmetric antibody trimer and a head-to-head interface between antibody trimers related by 2-fold symmetry that combined to yield a higher-order complex comprising two trimeric layers of antibodies (Figs. 6-7)
- Structure-function studies reveal a new antibody-based structural mechanism with dual homotypic interfaces mediating protection against varying numbers and spacing of minor repeats and major repeats.

## INTRODUCTION

Malaria presents a substantial burden on global human public health, causing over 263,000,000 cases and 597,000 deaths in 2023^1^. The majority of deaths are in children under 5 years of age and result from *Plasmodium falciparum* infection, with morbidity and mortality concentrated in Africa. There are a number of tools such as vaccines, mosquito control measures and preventive drug treatments that are used to control malaria. Monoclonal antibodies are a potential new intervention to prevent malaria infection and transmission^2–4^. The circumsporozoite protein (CSP) is presented at high density on the surface of the sporozoite stage of the parasite life cycle and represents an important target for malaria prevention by monoclonal antibodies and vaccines^5–7^. Monoclonal antibodies against CSP are safe and mediate high-level protection against malaria in a number of recent Phase 2 studies in Africa^8–10^. While these studies highlight the potential of monoclonal antibodies to control malaria, a key driver for their broad implementation will be to minimize their cost. Thus, active investigation is underway to generate more potent antibodies.

CSP is mostly comprised of repeated peptide motifs and appears not to have a single unique structure, with recognizing antibody inducing different conformations. In *P. falciparum* CSP, the dominant peptide repeats consist of an Asn-Ala-Asn-Pro (NANP) tetrapeptide for the major repeats and an Asn-Val-Asp-Pro (NVDP) for the minor repeating motifs. Most *P. falciparum* strains, including the widely used 3D7 strain, contain three closely spaced minor repeats within the CSP central repeat region. Each minor repeat is commonly separated by a single major repeat, and CSP often has another distal minor repeat after a dozen or so major repeats. The L9 monoclonal antibody preferentially targets the cluster of three NVDP-minor repeats on *P. falciparum* CSP, utilizing an asymmetric trimer^11,12^. Thus, L9 mediates protection primarily through high-affinity interactions against the minor repeat. Other potent monoclonal antibodies in clinical development against PfCSP target the major repeat^13–15^ or the junction region^5,16–18^.

*P. falciparum* has polymorphisms in the *Pfcsp* gene, including variation in the numbers of major and minor repeats, as well as the spacing between them. Studies have shown that L9’s protective mechanism is reduced when only a single minor repeat is present, and thus L9 requires at least two minor repeats in the PfCSP molecule for full protection^19^. Structural studies also suggest that the L9 homotypic structures optimally recognize minor repeats that are in close proximity^11^; still, L9 is substantially protective against strains with two spatially separated minor repeats. Very few strains have no NVDP repeats, and around 8.4% of strains analyzed in a recent study (32/382 strains registered in GenBank in 2022, which may not reflect actual field distributions) had ≤1 NVDP repeat, against which L9 would thus be less effective^11^. Thus, a modified version of L9 that could recognize rare strains with only a single minor repeat could enhance breadth across all strains.

Antibody engineering is a useful tool to improve antibody breadth and potency in other systems. Our prior efforts showed that directed evolution via yeast display could be useful to improve antibodies for enhanced performance against infectious diseases, including for antibodies targeting the fusion peptide of HIV-1^20,21^ and the junctional peptide motif of PfCSP^16^. In this study, we hypothesized that systematic mutagenesis and high-throughput screening could be used to evolve versions of L9 with sufficient affinity to recognize strains with minor repeat variations compared to the benchmark 3D7 strain. Such an evolved variant would have sufficient breadth to protect against malaria variants with potential resistance to the L9 antibody.

Engineering the L9 antibody via yeast display, we found that several mutations on L9 that could improve affinity to PfCSP peptides which encompassed minor and major motifs, and one of the resulting variants provided improved protection against transgenic *Plasmodium* strains with one or no minor repeats. Structural and sequence analysis revealed this variant to have increased affinity to the major repeats, and alterations that introduced a second homotypic interaction leading to a head-to-head dimerization. These data reveal a new structural pathway through which to develop effective anti-malaria biologics against the repeat regions of *P. falciparum* CSP.

## RESULTS

### Directed evolution to improve L9 affinity against selected PfCSP antigens

L9 binds to PfCSP predominantly through the NVDP minor repeat, with lower affinity contacts to the NANP major repeat that are not sufficient for full protection (**Fig. 1A**)^19^. We sought to further improve L9’s affinity to its targeted epitopes, selecting first for affinity to Peptide 22, the sequence motif used to define L9 specificity which contains 2 minor repeats and 2 major repeats. We also selected libraries for maintained binding to NTDS_5/3, a truncated version of the PfCSP molecule that could present the same peptide antigens in a more native context as probes (**Fig. 1B**)^16^. We employed a stepwise approach to generate L9-variant libraries, using several rounds of yeast display screening to progressively enrich for variants with improved affinity to Peptide 22, and also for preserved binding to NTDS_5/3 (**Figure 2A**). First, we generated site-saturation mutagenesis (SSM) libraries encompassing every possible single amino acid mutation on the heavy (VH) and light (VL) chains of L9^22^. The SSM libraries were cloned into yeast surface display and labeled for FACS with Peptide 22 (Pep22, 0.5 nM), which contains two major NANP and two minor NVDP motifs. Sorted L9-SSM libraries were analyzed by flow cytometry, with improved affinity validated using a high affinity gate that measured the ratio of Pep22 binding (Geomean _PE_) to Fab display (Geomean _FITC_). After three rounds of affinity-based FACS, the enriched populations had a higher mean fluorescence intensity (MFI) compared with the Pre-sort libraries and wild type (L9_WT), including for binding to peptide antigens and the NTDS_5/3 truncated PfCSP (**Figure 2B, Figures S1-S3**).

**Figure 1.**
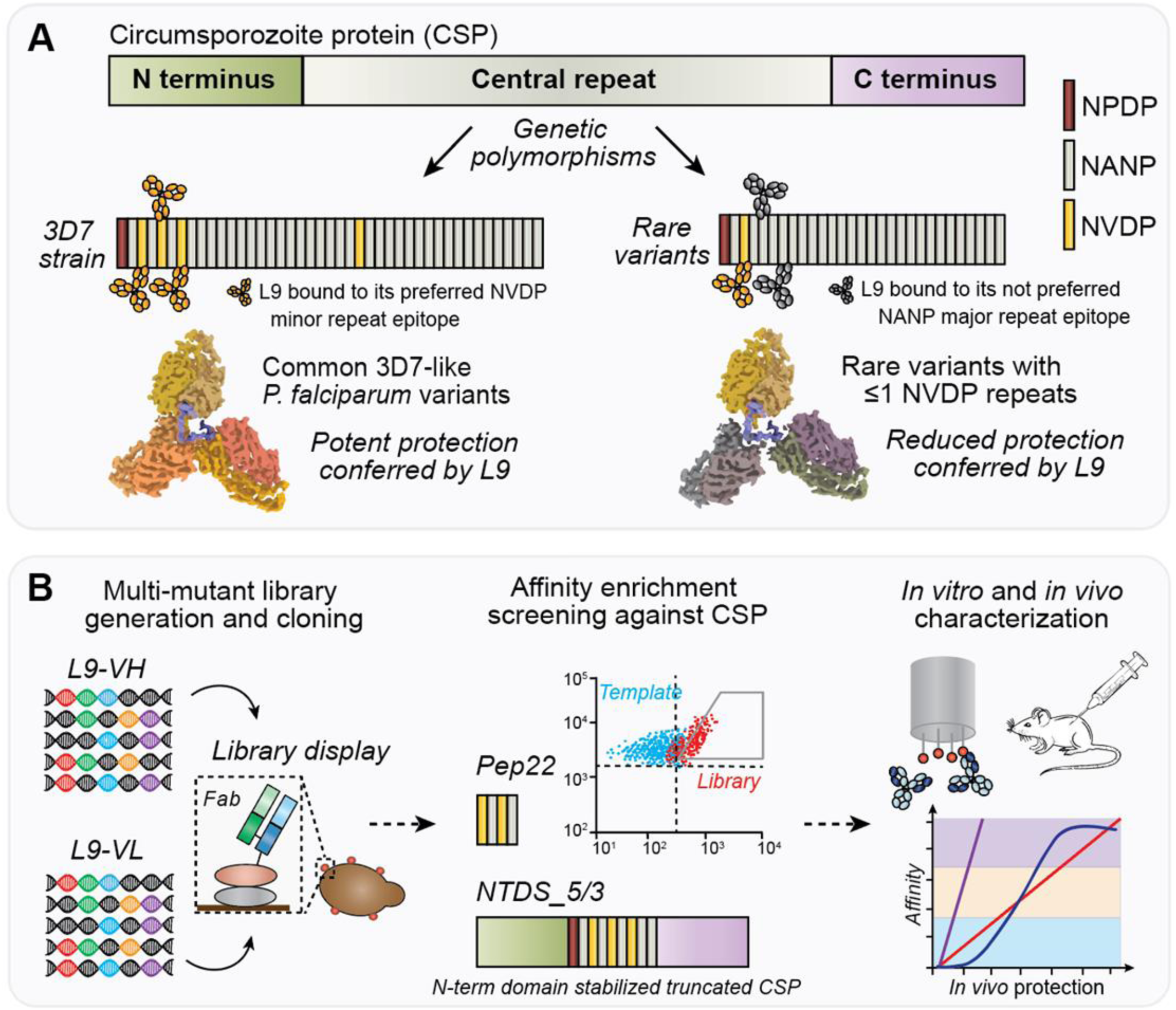
Improving affinity to target the CSP minor repeat enhances L9 antibody protective breadth. **(A)** The CSP minor repeat (NVDP) minor repeat is the principal binding target of the potent anti-malarial antibody L9. Genetic polymorphisms are present among *P. falciparum* strains in the number and spacing of NVDP motifs contained in the CSP repeat regions, and the loss of NPNV epitopes reduces the *in vivo* protective potency of the L9 antibody. **(B)** To enhance L9 affinity against the CSP antigen, multi-mutation libraries were generated from the L9 template and selectively screened against CSP peptide repeat motifs using yeast display. Libraries were first selected against Peptide22 (Pep22), which contains two minor and two major repeats, and subsequently selected against NTDS_5/3, a truncated CSP molecule that presents peptides in a more native-like conformation. Identified L9 variants were characterized for binding affinity *in vitro* and for protective potency *in vivo* against a panel of malaria antigens.

**Figure 2.**
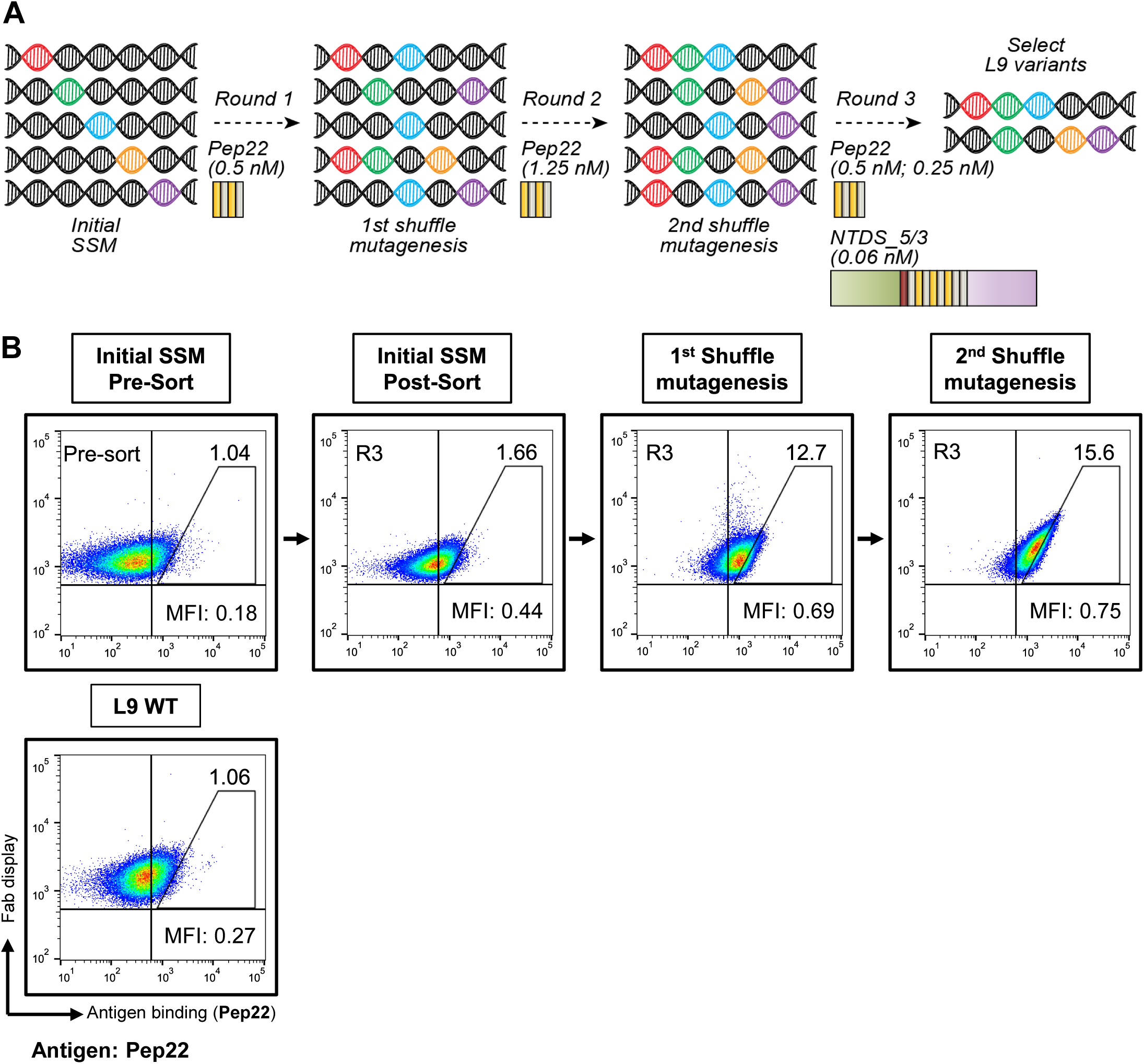
Library generation and yeast display screening evolves new L9 variants with improved affinity to peptide 22. **(A)** The L9 antibody was used as template for multiple rounds of directed evolution for affinity improvement against malaria antigens. First, site-saturation mutagenesis (SSM) and sorting was used against 0.5 nM Pep22 (2 NVDPs). Next, enriched variants further diversified using SSM and DNA shuffling, followed by sorting against 1.25 nM Pep22. Finally, the enriched libraries were shuffled again and sorted under increasingly stringent conditions by progressively lowering Pep22 concentrations during FACS, with a final enrichment step against NTDS_5/3 (a truncated CSP with 5 NANP / 3 NVDP repeats). **(B)** Representative flow cytometry plots across the screening experiments. Each population was labeled with 62.5 pM Pep22. A high-affinity gate is drawn for comparative purposes, and the percentage of the high affinity population is shown in the upper-right quadrant. The mean fluorescence intensity ratio (MFI) is calculated as (Geomean _PE_ / Geomean _FITC_) to quantitatively represent the normalized affinity of Fab molecules. See also Figs. S1-3.

Next, we performed two rounds of shuffling mutagenesis to the enriched population of single mutation variants, combining another round of SSM along with DNA shuffling to generate multi-mutant libraries^23,24^. Yeast display libraries of the 1^st^ shuffle mutagenesis were labeled with Pep22 (1.25 nM) for FACS. After the 1^st^ shuffle mutagenesis, the enriched population showed at least 4-fold higher MFI than Pre-sort populations. Libraries were enriched against Pep22 such that majority of the Fab-displaying population bound to Pep22 and NTDS_5/3 even when analyzed with antigen at a low concentration of 62.5 pM, where much of the wild-type population was not binding antigen (**Figure 2B, Figures S2-S3**). We iterated the shuffle mutagenesis once more to further identify multi-mutation L9 variants. Due to the very large potential library sizes that can be achieved by DNA shuffle mutagenesis^25^ we first performed Pep22 MACS to remove non-expressing Fab clones prior to affinity enrichment by FACS. We also lowered the Pep22-labeling concentration again and screened the libraries using NTDS_5/3 in the last round of FACS sorting to ensure that selected variants would bind the major and/or minor repeat peptides when presented in the context of a CSP molecule (**Figure 2A**). The libraries were nearly completely enriched with Pep22-binding populations and also showed a higher MFI against NTDS_5/3 than L9_WT, which suggested that the sorted libraries were enhanced in their affinity against both Pep22 and PfCSP compared to that of the already high-affinity L9_WT molecule (**Figures S2-S3**).

### Bioinformatic mining of affinity-enhanced L9 variants

We assessed the genetic diversity of enriched libraries across each round of screen by next-generation sequencing (NGS). L9 variants were selected for *in vitro* characterization based on the prevalence of a given mutation or sequence in enriched library populations and also based on the enrichment ratio (ER) of L9 variants in the last round of screening. Most libraries were sequenced using the Illumina MiSeq platform. For the last round of libraries, we used the Pacific Biosciences Sequel II platform that offers longer read length for complete paired VH:VL sequences, albeit with lower read counts than Illumina MiSeq. We combined the Pacific Biosciences data with Illumina MiSeq data for higher throughput of separate heavy and light chains^26^. Due to higher insertions/deletion error rates at homopolymer regions in Pacific Biosciences data, we cross-checked all selected sequences by evaluating the separate VH and VL sequences from Illumina data using the same criteria, and we identified 36 paired VH:VL sequences for expression. There were 32 different VH mutations and 15 different VL mutations present in the selected variants, among which 19 VH mutations and 3 VL mutations were observed only in the last round of DNA shuffling and screening, including 5 unique single amino acid mutations; VH_S25D, VH_T69E, VH_D73R, VL_F80_H_W, and VL_G57_L_H (template numbering, see **Table S1**). At the single amino acid level, six more mutations (VH_V50A, VH_I58T, VH_M81L, VH_G103R, VL_Q27R, and VL_Q89A) consistently appeared to be most prevalent from the variants screened in DNA shuffling rounds. Interestingly, one VH_F53G mutation was strongly enriched in the first step of DNA shuffling to being contained in 58.7% of sequences, then reduced to 1.8% after the second round of shuffling, which implied that it conferred superior properties to L9_WT but was out-competed in the later screening rounds (**Figure 3A, 3B**). While some of the most prevalent mutations occurred in complementarity determining regions (CDRs), the mutations identified were not limited to the CDRs (**Figure 3**). Because most individual mutations were already showing significant enrichment after the first round of DNA shuffling, the most prevalent mutations going into later screening rounds did not show strong enrichment (**Figure 3A, 3B**). Instead, the later rounds selected for optimal combinations of amino acid mutations and depleted the libraries of mutations that provided an advantage over wild-type L9 but that were outcompeted by other clones, such as VH_F53G. The selected enriched variants after three rounds of directed evolution were then expressed as soluble antibodies for characterization.

**Figure 3.**
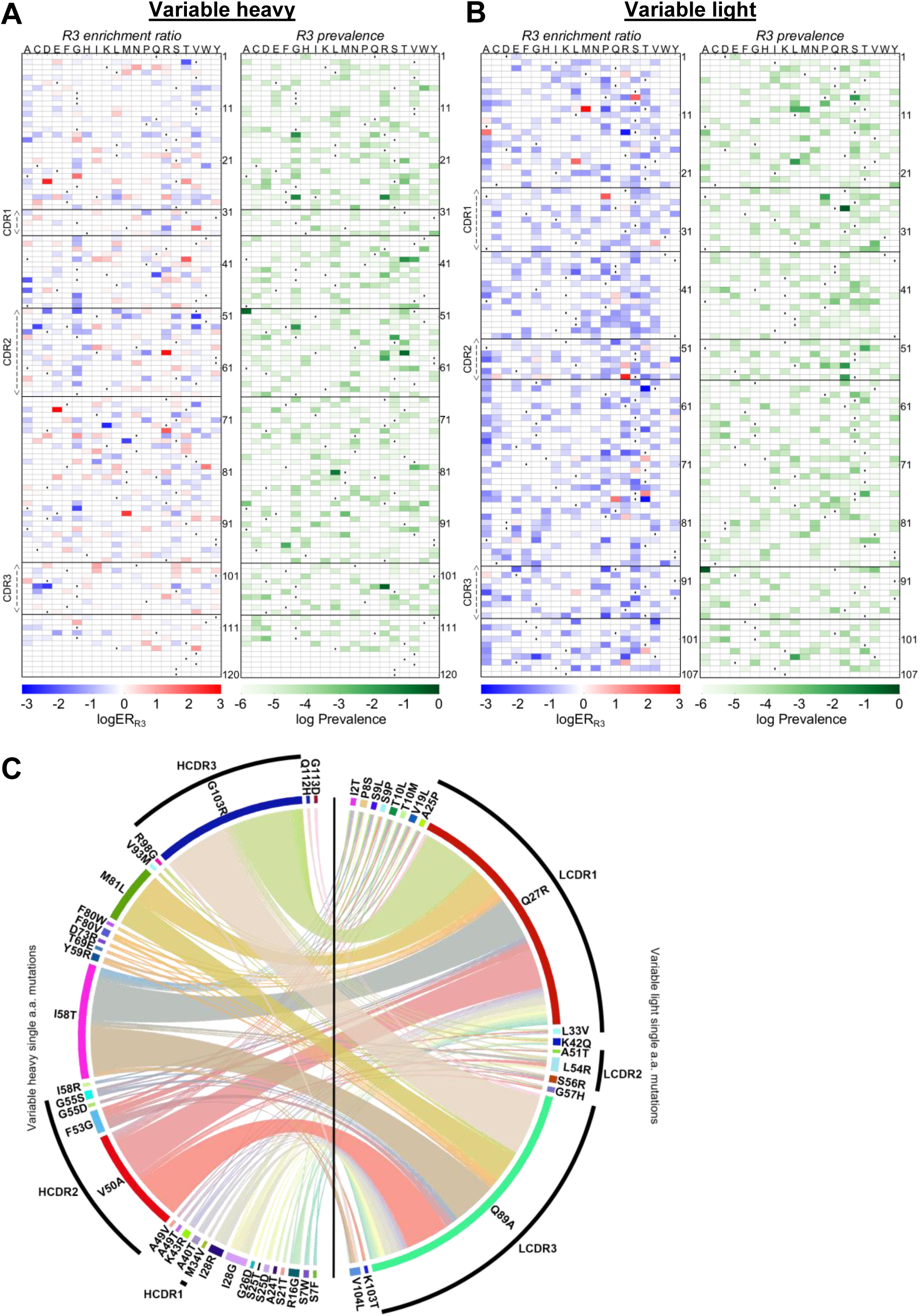
Bioinformatic analysis reveals affinity-enhanced VH:VL sequences after Round 3 (R3) of the 2^nd^ round of shuffle mutagenesis. The round 3 (R3) enrichment ratio (left) and prevalence (right) for amino acid mutations identified from **(A)** VH sequences and **(B)** VL sequences. Wild-type amino acids are indicated with dots (·). **(C)** Mutational pairings between single amino acid mutations identified from VH (left half) and VL (right half) sequences. The amino acid residues followed template numbering. See also Fig. S4.

### Binding analysis and *in vivo* protection by L9 variants

We first tested the 36 selected L9-variant antibodies using biolayer interferometry (BLI) to quantify affinity improvements (**Figure 4A, Table S1**). We found that nearly all the selected variants had improved affinity against Peptide 22, with one variant (yd01) showing enhanced affinity by approximately 100-fold. Nearly all variants maintained binding affinity to full length PfCSP, although only a select few variants showed improved affinity to the benchmark 3D7 strain PfCSP.

**Figure 4.**
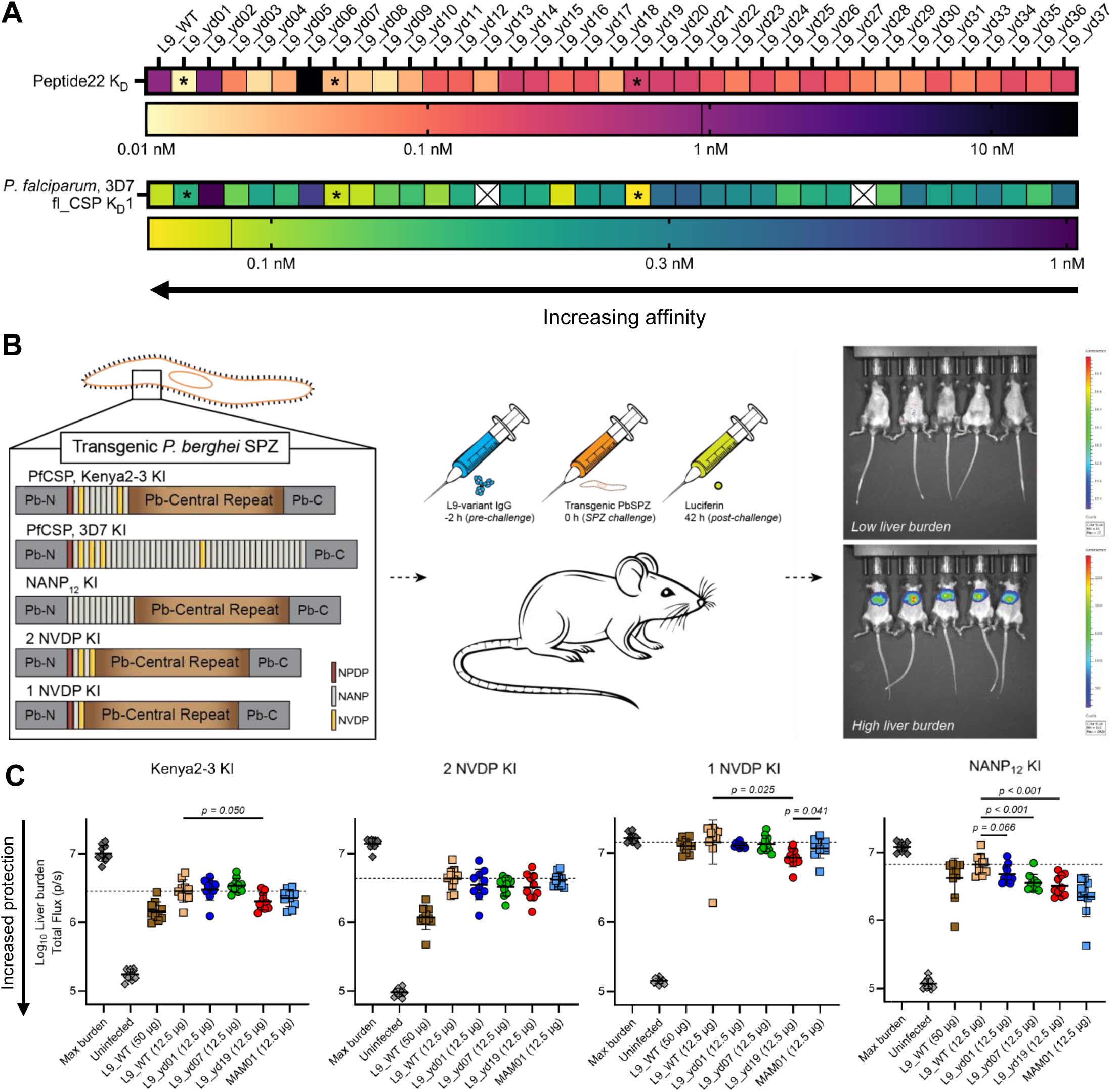
*In vitro* and *in vivo* characterization reveals functional improvement for affinity-enhanced L9 variants against multiple transgenic strains, including the Kenya2-3 KI strain and a major repeat-only transgenic. **(A)** Biolayer interferometry (BLI) analysis against Peptide 22 (containing 2 NVDPs) and full-length CSP (fl_CSP) of *P. falciparum* 3D7. L9_WT affinity is marked with a vertical line on the legend bar. L9 variants marked with an asterisk (*) were selected for detailed *in vivo* characterization. **(B)** In the transgenic sporozoite intravenous (IV) challenge model, mice were administered IgG from L9 variants or control antibodies (−2 h), followed by IV challenge with *P. berghei* sporozoites (PbSPZ) encoding different PfCSP knock-in sequences (0 h), and subsequent luciferin-based evaluation (42 h). Each group comprised n=10 mice, with animals receiving the indicated antibody dose. **(C)** Transgenic PbSPZ challenge. Liver infection burden was quantified, and data are shown as mean ± standard deviation; mean liver burden total flux for L9_WT at 12.5 µg/mouse is indicated by a dashed line. Statistical significance between L9_WT and each L9 variant group at 12.5 µg were determined using the ordinary one-way ANOVA test with a two-tailed P value calculation and a Dunnett’s multiple test correction. Statistical significance was also analyzed between L9_yd19 and MAM01 using an unpaired t test with a two-tailed P value calculation; the two groups were significantly different when tested against 1 NVDP KI PbSPZ. See also Table S1 and Fig. S5.

Based on these affinity data, we selected a panel of seven of the improved variants for *in vivo* challenge analysis. In these models, the CSP protein of a *P. berghei* strain expressing luciferase had been replaced with a transgenic *P. falciparum* CSP. Mice were challenged intravenously with the transgenic PfCSP *P. berghei* sporozoites labeled with luciferase, and parasite burden in the liver was quantified^16,19,27^. For our initial analysis we used a strain which has only two widely separated NVDP-minor repeats that is similar to an isolate from Kenya, termed Kenya2-3 knock-in, as well as the more commonly used benchmark 3D7 strain that contains the standard complement of three closely spaced NVDP-minor repeats, as well as an additional minor repeat in the central repeat region (**Fig. 4B**). With a 25 microgram/mouse antibody dose, L9 variants were comparable to L9 in mediating reduction of liver burden against the Kenya2-3 or 3D7 knock-in strain (**Figure S5**).

We selected three variants (yd01, yd07, and yd19) for broader characterization against a panel of transgenic strains based on their ∼1-log reduction in liver burden in 25 microgram dose assays, and for their diversity of affinity phenotypes. yd01 had the highest Peptide 22 affinity and lower CSP affinity than WT; yd07 had moderately improved affinity to both Peptide 22 and CSP; and yd19 had some improved affinity to Peptide 22 and the very highest improvement against CSP (**Table S1**). These three variants all showed *in vivo* protection against the Kenya2-3 KI strain at 12.5 microgram/mouse, with a modest yet statistically significant improvement for L9_yd19 compared to L9 (**Fig. 4C**). As an additional comparison, MAM01, an antibody with high affinity binding to major repeats, showed comparable reduction of liver burden at the same dose.

We next sought to understand how the reduction in liver burden by yd19 and the other selected variants was affected by changes in the number of NVDP repeats. The Kenya 2-3 strain has two separated NVDP motifs. Thus, we also tested the antibody variants against transgenic chimeric parasites with 2 NVDP motifs with close spacing and also with just a single NVDP. While there were no significant differences in liver burden against the 2 NVDP strain by the L9 variants, L9_yd19 showed the lowest liver burden amongst all variants tested against the 1 NVDP strain (p=0.025 vs. L9_WT) and also compared to MAM01 (p=0.041), indicating that the mutations in yd19 had overcome the substantial loss in potency known to occur in parasites with only a single NVDP motif (**Figure 4C**). Finally, we analyzed the performance of all three strains against a KI strain with NANP_12_, which contains only NANP motifs and no NVDP motifs. L9_yd19 showed significantly reduced liver burden against the NANP_12_ KI strain compared to L9_WT (p<0.001), with no statistically significant difference compared to that of the malaria-protective antibody MAM01. which binds preferentially to the major repeat (**Figure 4C**)^13^. These *in vivo* challenge data demonstrated that the mutations selected by yeast display were effective in enhancing L9 affinity, and also in broadening its targeted epitope to include both the NVDP minor repeat, as well as the NANP major repeat regions.

### Functional correlates of antigen affinity with protective potency across malaria knock-in strains

We sought to understand any functional and affinity-based correlates of protection for the knock-in strains used. In these affinity assays we evaluated L9_WT and the three variants yd01, yd07, and yd19 (selected across a broad affinity range for detailed testing in challenge studies). We used a broad panel of CSP antigens that included PfCSP_Kenya2-3, NTDS_5/3, PfCSP_3D7 (**Figure 5, Table S2**), and also the antigens Peptide 22 (major and minor repeat), Peptide 21 (junction region, major repeat, minor repeat), and Peptide 29 (major repeat only) (**Figure S6, Table S2**). We found that protection against 2 NVDP KI, 1 NVDP KI, and NANP KI strains was correlated with affinity to the Kenya2-3 strain (p<0.02 in all cases), and also with K_D_2 of PfCSP_3D7 with high R^2^, but p>0.05. Interestingly, protection against these strains was not as closely correlated with affinity to NTDS_5/3, or with K_D_1 of PfCSP_3D7 (**Figure 5, Table S2**). Very few peptide affinity correlates were significant with the exception of Peptide 29 affinity correlated with NANP12 protection (p=0.045), although several correlations had strong R^2^ values (**Figure S6, Table S2**). Neither peptide affinities nor CSP affinities were effectively correlated with protection against the Kenya2-3 KI strain (**Figure S7, Table S2**).

**Figure 5.**
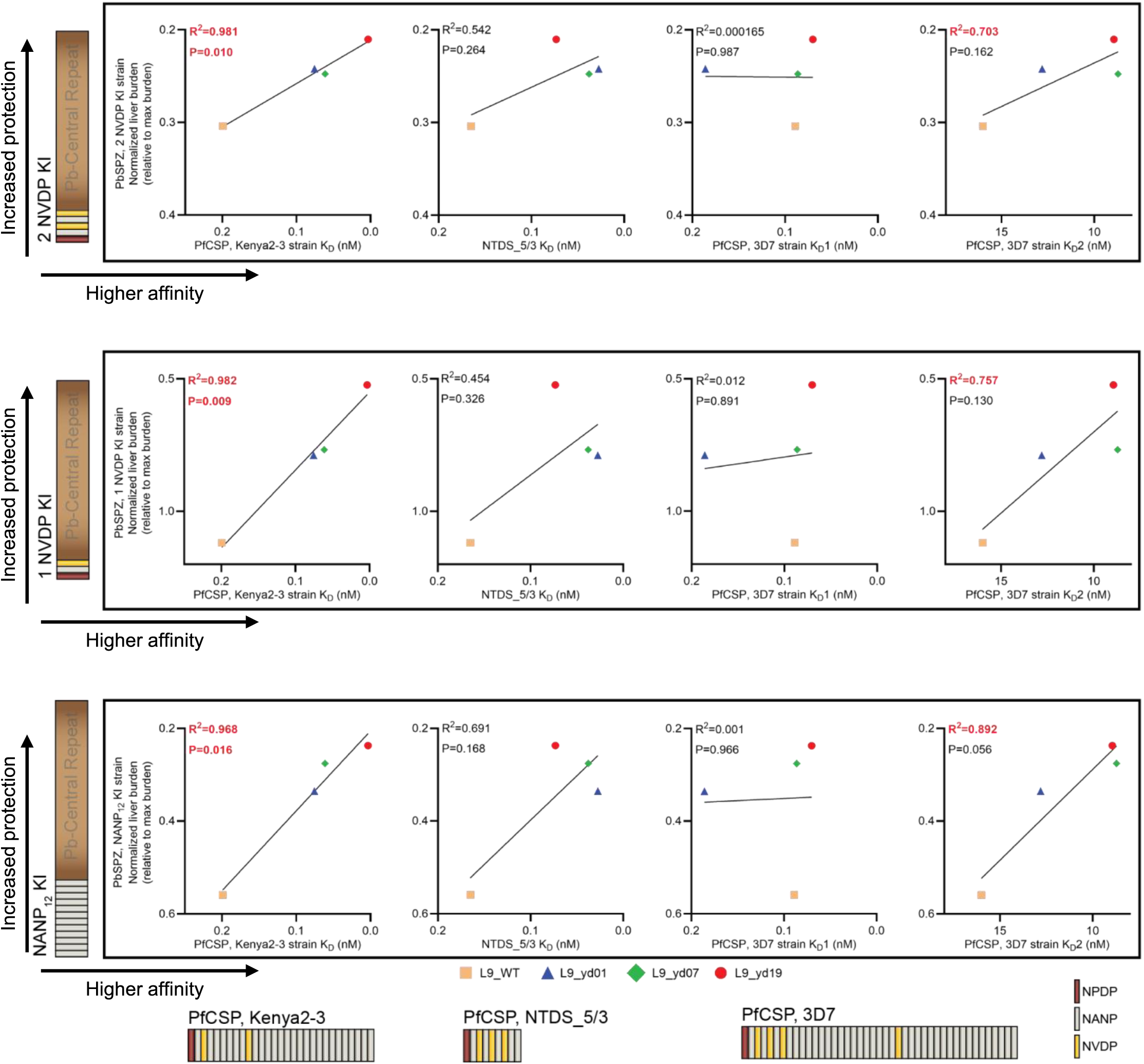
Affinity correlations reveal that improved protection breadth correlates with affinity to multiple malaria antigens. Correlation between binding affinity to PfCSP Kenya2-3 strain, NTDS_5/3 or PfCSP 3D7 strain (x-axis, linear scale) and normalized liver burden against various transgenic PbSPZs (y-axis, linear scale). Pearson’s correlation coefficient (R) was calculated using the median normalized liver burden values, with corresponding two-tailed P-values and 95% confidence intervals. A linear regression was fit based on a Pearson’s correlation analysis. See also Fig. S6.

### Cryo-EM structures for L9_yd19 in complexes with PfCSP antigens

To gain mechanistic insights into the enhanced protective efficacy of L9_yd19 against Kenya2-3 transgenic strain, we sought to obtain its cryoEM structure in complex with PfCSP from both 3D7 and Kenya2-3 strain. L9_yd19 antigen binding fragment (Fab) was complexed with PfCSP (3D7 or Kenya2-3) at a 2:1 molar ratio and single particle cryoEM data was collected.

The 2D classification of L9_yd19:PfCSP-3D7 showed a major class consisting of three L9_yd19 Fabs bound to PfCSP-3D7 (**Figure 6A**, upper left panel), similar to L9_WT in complex with PfCSP (3D7)^11,12^. A small number of classes were also observed that showed more than three L9_yd19 Fabs bound to PfCSP-3D7, however, a high-resolution structure could not be obtained because of low overall number of particles. The structure of three Fabs bound complex was obtained at a resolution of 3.4 Å (**Figure 6A**, upper right panel). The density for 27 residues of PfCSP-3D7 was observed and the rest of the CSP was disordered. L9_yd19 had five mutations relative to L9_WT; two in heavy chain (V50A, I57T) and three in light chain (Q27R, Q89A, K103T) (**Figure S8**). The V50 in CDRH2 and Q89 in CDRL3 surround the CDRL3 loop, which is critical for the interactions of L9’s preferred epitope (NPNV) with the light chain. The mutation of both residues to smaller residues (V50A, Q89A) may provide additional flexibility to the CDRL3 loop for a more favorable interaction with the NPNV epitope (**Figure 6A**, lower panel).

**Figure 6.**
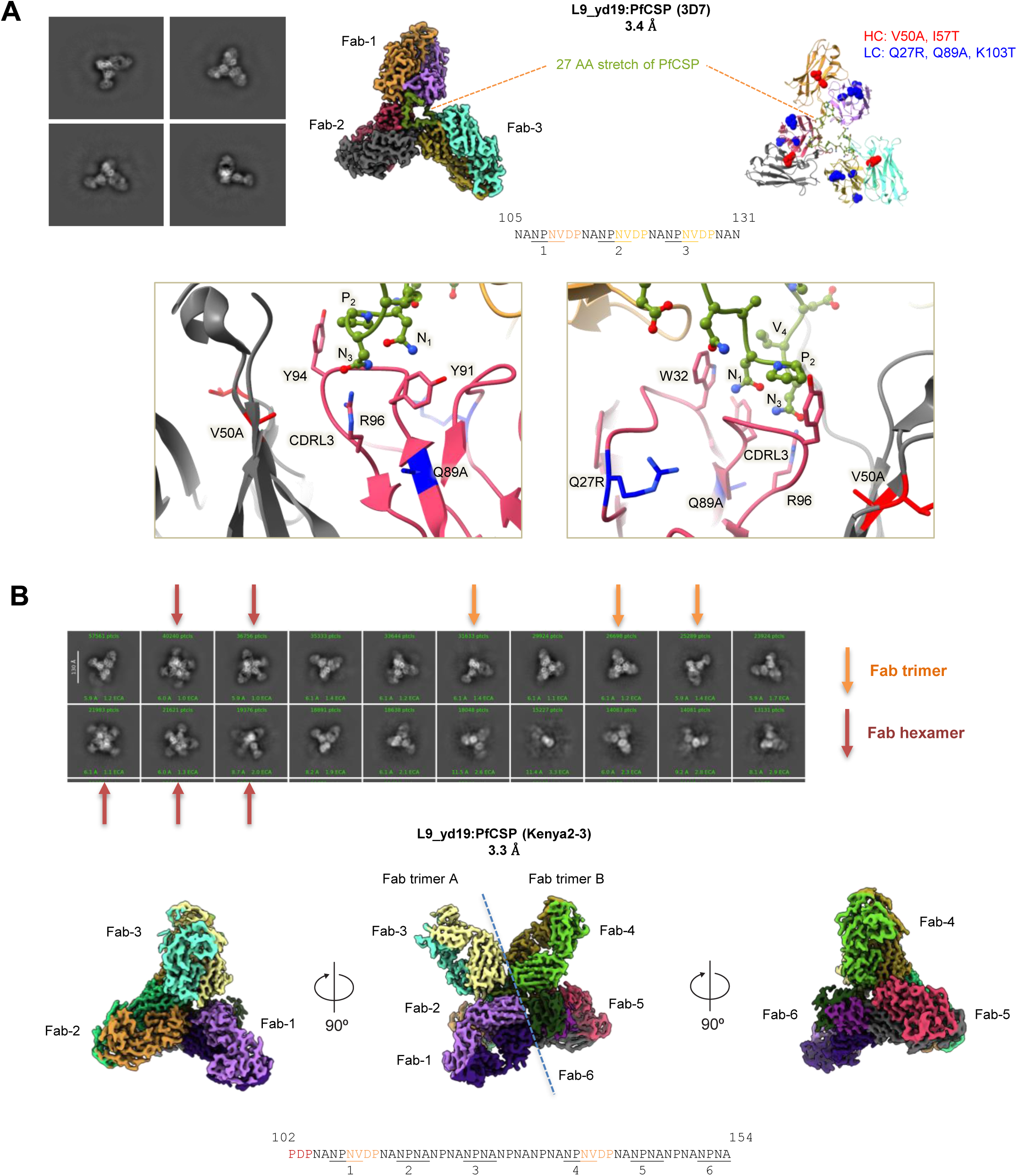
Structural analyses of L9_yd19 and PfCSP (3D7 or Kenya2-3) reveals a new interaction mode. (**A**) CryoEM structure of L9_yd19 in complex with PfCSP (3D7) at 3.4 Å resolution. Heavy chain mutations are shown in red and light chain mutations are shown in blue. 27 residue stretch of PfCSP is ordered and the rest of the PfCSP is disordered. (**B**) CryoEM structure of L9_yd19 in complex with PfCSP (Kenya2-3) at 3.3 Å resolution. 2D classes show both three L9_yd19 Fabs (orange arrow) and six L9_yd19 Fabs (red arrow) bound to PfCSP (Kenya2-3). 3D reconstruction for six L9_yd19 Fabs bound to PfCSP (Kenya2-3) is shown.

The 2D classification of L9_yd19 in complex with PfCSP-Kenya2-3 showed an alternate binding mode with the majority of the classes containing more than the traditional three Fabs bound; classes with three Fabs bound were also observed (**Figure 6B**, upper panel). The 3D reconstruction yielded a 3.3 Å resolution map with six L9_yd19 Fabs bound to a 53-residue stretch of PfCSP-Kenya2-3. Since the preferred high affinity epitope (NPNV) is separated by 6 NANPs in PfCSP-Kenya2-3, this higher order complex utilized the lower affinity epitope (NPNA) to form two layers of Fab trimers stacked on top of each other (**Figure 6B** bottom panel), one layer recognizing minor-major-major repeats and the other layer recognizing another sequence of minor-major-major repeats. This unique binding modality of L9_yd19 thus provided additional Fab-Fab homotypic interactions that enabled enhanced affinity and protective efficacy against the Kenya2-3 transgenic sporozoite. Overall, while L9_yd19 displayed a canonical three Fab binding mode with PfCSP-3D7, it also showed a new binding mode in which six L9-yd19 Fabs bound to a combination of minor and major repeats.

### L9_yd19 engineered mutations enable new Fab-Fab homotypic interactions

To investigate the critical residues in L9_yd19 that enabled additional homotypic interactions, we analyzed the structural differences between the canonical three Fab complex (3D7) versus the newly observed six Fab-CSP complex (Kenya2-3) (**Figure 7A**). The analyses revealed Fab-1 (numbered based on distance from the N-term of CSP) to bind the preferred high affinity epitope (NPNV), anchoring the complex. The subsequent Fabs (2 and 3) bound to a secondary low-affinity epitope (NPNA) separated by a four-residue linker (D/NPNA) and potentially stabilized through side-to-side Fab-Fab homotypic interactions in the atypical trimer context. These interactions utilize homotypic contacts between heavy chain of one Fab to the light chain of an adjacent Fab (**Figure 7B**, upper left panel).

**Figure 7.**
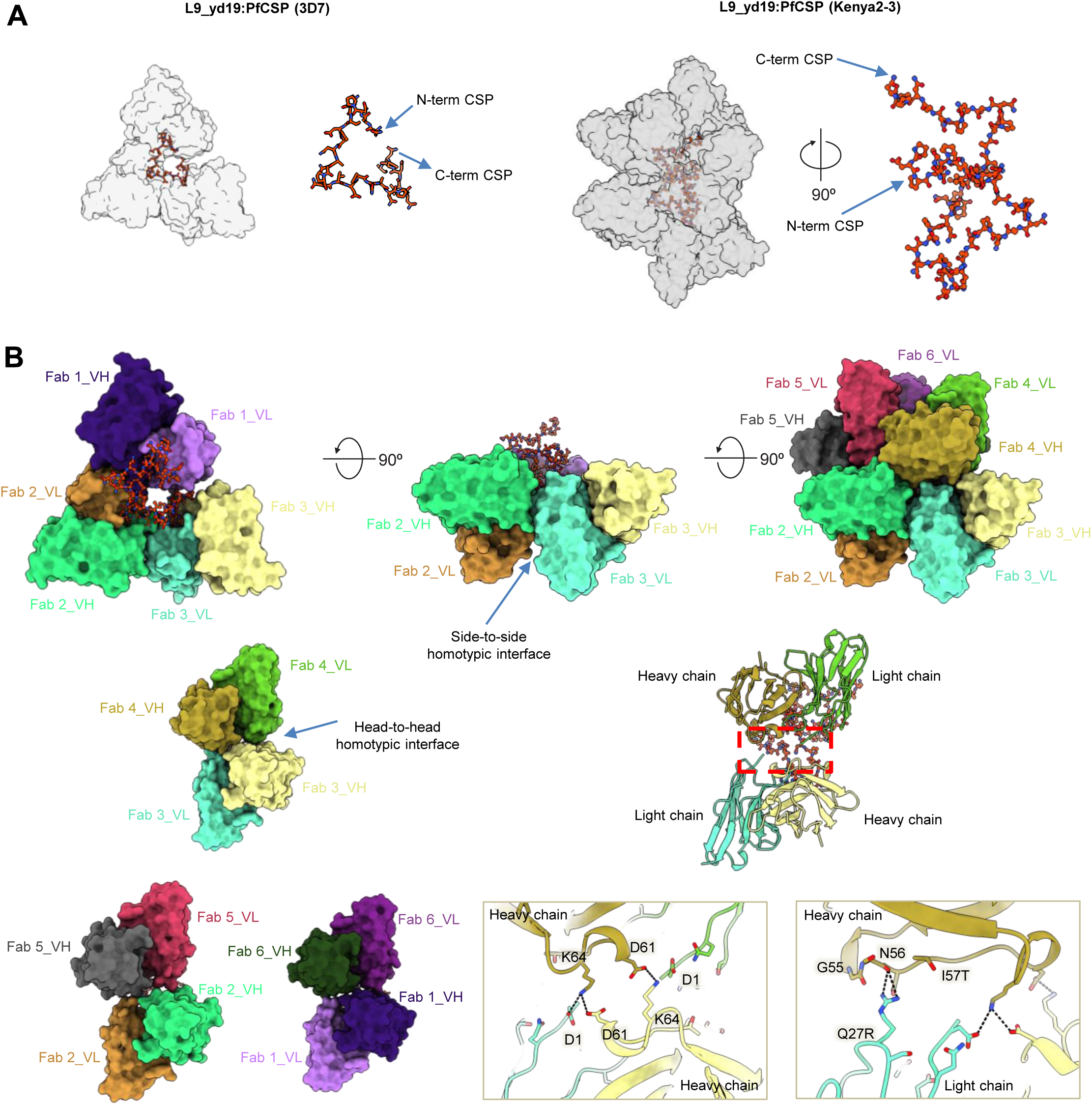
L9_yd19 engineered mutations stabilize a new head-to-head homotypic interface allowing up to six Fabs to bind to PfCSP (Kenya2-3). **(A)** Structural comparison of L9_yd19 bound to PfCSP-3D7 and PfCSP-Kenya2-3. PfCSP is shown in red sticks with the N- and C-termini labeled. (**B**) L9_yd19 in complex with PfCSP-Kenya2-3. Fabs (3,4), (2,5), and (1,6) are related by 2-fold (C2) symmetry. Critical residues generating the head-to-head homotypic interface are shown in insets.

The engineered mutations in L9_yd19, specifically VL_Q27R and VH_I57T, facilitated a second layer of Fab trimer comprising of three Fabs (4-6) to stack on top of the first layer of Fab trimer (1-3). The Fab-3 and Fab-4 were separated by an eight-residue linker (NPNPNPNA), and this arrangement allowed Fab-4 to bind the primary high-affinity epitope (NPNV) and act as an anchor for the upper layer of Fab trimer. In the upper layer, Fabs (4-6) interacted in the same fashion as the bottom layer trimer. A unique head-to-head Fab-Fab interaction was observed between the Fabs from the upper and the lower layer (**Figure 7B**, top right panel). This novel arrangement allowed additional inter-Fab contacts and stabilized the higher order complex.

Symmetry analysis revealed that the upper layer Fabs exhibited a two-fold (C2) symmetry with the lower layer Fabs (Fab-4 and −3, Fab-5 and −2, Fab-6 and −1) (**Figure 7B**, lower left panel). This head-to-head homotypic interface was stabilized by polar interactions between both the heavy and light chains of the two Fabs. VL_D1 and VH_D61 from Fab-3 formed a reciprocal salt bridge with VH_K64 on Fab-4, which created a symmetric pair of salt-bridges between Fabs 3 and 4 and further stabilized the interface. The introduced mutations in L9_yd19 (Q27R_L_ and I57T_H_) also stabilized this interface via hydrogen bond interactions between VL_Q27R and the backbone carbonyl of VH_N56 and a polar interaction between VL-Q27R and VH_I57T (**Figure 7B**, lower right panel). Similar set of interactions are observed between Fabs (2 and 5), and Fabs (1 and 6).

While structural analysis defined location of Fabs, the location of the connecting constant regions needed to be inferred using geometric constraints. The three Fabs making up each recognition layers had C-terminal portions of the Fab-heavy chain (C-terminus of CH1) that ranged in distance from 93-107 Å, likely too long to be connected by a standard IgG1 constant region^11,12^, but potentially by an IgG3^28^. Meanwhile, CH1-CH1 distances for Fabs 3-4, 2-5, and 1-6 ranged from 106-108 Å, also too long for an IgG1, but potentially for an IgG3. In any case, each of the individual Fabs of the L9_yd19 recognition complex, could also be contributed by separate IgGs.

Altogether, these findings reveal how L9_yd19 circumvents the spatial requirements of L9_WT, which requires two high-affinity epitopes (NPNV) to be separated by a four-residue linker for optimal potency, via the formation of higher order complexes through head-to-head homotypic interactions.

## DISCUSSION

Malaria continues to be a global threat to public health, and efficient, low-cost interventions are urgently needed to complement existing control measures. Here, using yeast display and site-saturation mutagenesis, we engineered the potent L9 antibody variant, L9_yd19 for improved affinity against CSP. We found that L9_yd19 enhanced affinity to the NANP major repeat compared with L9_WT, while also showing enhanced affinity to the NVDP minor repeat. L9_yd19 mediated reduction in liver burden with chimeric transgenic parasites containing only NANP repeats, which may expand breadth against rare strains with a single NVDP minor repeat or improve potency against strains with two minor repeats that are spaced.

The ability of L9_yd19 to recognize the NANP epitope appears to result from a unique binding mode that relies on a new form of head-to-head homotypic Fab-Fab interactions. These structural mechanisms use previously unreported Fab-Fab interactions with the parental antibody L9, to “anchor” multiple trimeric Fab complexes to CSP, either using one minor repeat epitope with two subsequent major repeat epitopes, or using entirely major repeat epitopes as in the protective efficacy of L9_yd19 against the NANP_12_ KI strain. The engineered mutations in L9_yd19 promote the formation of a higher order antigen-antibody complex via stabilization of the head-to-head homotypic interface by polar interactions. This unique recognition mode, with two distinct homotypic interfaces, enables L9_yd19 to achieve enhanced protective efficacy against PfCSP strains that lack the optimal high-affinity minor repeat epitope or preferred spacing.

We note that strains with fewer and/or more sparse NVDP epitopes are rare in sequenced malaria genotypes^11^. It may be significant that the L9_yd19 antibody can reduce liver burden in mouse models with only NANP repeats, or modestly reduce liver burden with chimeric transgenic strains with a single minor repeat, potentially allowing for expanded breadth across all strains. L9 variants utilizing the dual homotypic mechanism of recognition revealed by L9-yd19 could potentially be further improved by increasing major repeat recognition, or by utilizing IgG3-linkers instead of IgG1.

In conclusion, we show here that directed evolution via yeast display provided improved variants for a very high-affinity antibody, and unveiled a new structural pathway for anti-malarial antibodies with improved protection against *Plasmodium* expressing globally circulating PfCSP variants. The affinity-and structure-based correlates of protection identified here can be used to further accelerate progress in antibody engineering against global *P. falciparum* strains. These data provide a new window of opportunity for antibody development against PfCSP and introduce new broadly protective antibodies that could efficiently prevent new malaria cases around the globe.

## ACKNOWLEDGMENTS

We thank the New York Structural Biology Center for access to the Tycho^TM^ NT.6. We thank C. França for assistance with figures, and members of the Structural Biology Section and Structural Bioinformatics Core, Vaccine Research Center, for discussions and comments on the manuscript. This research was supported in part by the Intramural Research Program of the National Institutes of Health (NIH). The contributions of the NIH authors are considered Works of the United States Government. The findings and conclusions presented in this paper are those of the authors and do not necessarily reflect the views of the NIH or the U.S. Department of Health and Human Services. Funding was provided by NIH grants DP5OD023118, R01AI181684, and R01AI192975, the Mark and Lisa Schwartz AI/ML Initiative, the MIT Research Support Committee, the MIT Department of Chemical Engineering, and the Ragon Institute of MGH, MIT, and Harvard. This study used the Office of Cyber Infrastructure and Computational Biology High Performance Computing cluster at the National Institute of Allergy and Infectious Diseases, Bethesda, MD.

## AUTHOR CONTRIBUTIONS

Designed experiments: J.C., B.M., P.T., K.M., M.P., T.Z., A.H.I., F.Z., R.A.S., P.D.K, B.J.D. Performed experiments: J.C., B.M., P.T., Y.F-G., H.L., I-T.T., N.K.H., B.J.F. Analyzed the data: J.C., P.T., Y.F-G., K.M., A.S.F., G.A.L., B.J.D. Writing: J.C., P.T., P.D.K, B.J.D. Reviewing and editing: all authors.

## METHOD DETAILS

### Site-Saturation Mutagenesis Library Generation and Cloning

DNA libraries containing all possible single mutations to the L9 template antibody were constructed on the VH and VL (kappa) variable region genes using a previously reported one-pot site saturation mutagenesis (SSM) protoco^l16,20–22^. Pools of mutagenic primers containing degenerate single codons were synthesized to cover all possible amino acids at each residue. As previously reported, a single-strand nick was introduced one strand of the plasmid was digested, and the remaining single-stranded circular DNA was used as a template for codon-targeted site mutagenesis. After primer annealing, the second strand was generated to yield dsDNA plasmids ready for transformation of electrocompetent bacterial cells. *E. coli* libraries were transformed and colony counts were used to estimate the transformed library size. Plasmid DNA libraries were isolated using a Maxi-Prep kit (Zymo Research), and expression cassettes and the yeast display pCT plasmid backbone were transformed into the yeast strain AWY101 using a high-efficiency homologous recombination, as previously reported^16^. The resulting yeast libraries were passaged twice in SDCAA media (20 g/L dextrose, 6.7 g/L yeast nitrogen base, 5 g/L casamino acids, 8.6 g/L NaH_2_PO_4_.H_2_O, and 5.4 g/L Na_2_HPO_4_; SDCAA from Teknova) at 30°C and 225 rpm for 12 to 24 hrs to ensure that individual yeast cells maintain a single copy of the plasmid. These initial libraries expressed in yeast cells are referred to as the *pre-sort* libraries.

### Single-Mutation Yeast Display Library Screening via FACS

Expression was induced in yeast antibody libraries, and the yeast were stained with antigen for FACS as in prior studies^24–26^. Affinity evaluation and fluorescence-activated cell sorting (FACS) were performed using a SONY MA900 flow cytometer, as in prior reports^26,29^. Yeast display libraries were incubated in SGDCAA induction medium (2 g/L dextrose supplemented to 20 g/L D-(+)-galactose, 6.7 g/L yeast nitrogen base, 5 g/L casamino acids, 8.6 g/L NaH_2_PO_4_.H_2_O, and 5.4 g/L Na_2_HPO_4_; SGCAA from Teknova) at 20 °C and 225 rpm for 36 hrs to induce expression and surface display of Fabs. We refer to Fab-displaying yeast cells in a library as VL+ populations, because the FLAG tag is located on the same polypeptide as the VL region. Induced yeast libraries were washed twice with ice-cold staining buffer (1x PBS with 0.5% BSA and 2 mM EDTA) and labeled with an anti-FLAG FITC Clone M2 monoclonal antibody (F4049, Sigma-Aldrich), and with 0.5 nM biotinylated Peptide 22 (NANPNVDPNANPNVD)^7^ conjugated with streptavidin-PE (S21388, Invitrogen), toe enable VL+ population quantification and affinity evaluation, respectively. In the first round of FACS, 3E+07 yeast cells from pre-sort libraries were gated into high-affinity groups based on the ratio of Fab display to antigen binding, as described previously^25, 26^. High-affinity populations were collected for subsequent rounds of affinity enrichment. Sorted yeast cells were expanded in low-pH SDCAA (20 g/L dextrose, 6.7 g/L yeast nitrogen base, 5 g/L casamino acids, 10.4 g/L trisodium citrate, 7.4 g/L citric acid monohydrate, pH 4.5) at 30 °C and 225 rpm for 24-48 hrs, followed by two additional rounds of FACS with reduced input of 1E+07 yeast cells.

### Shuffled Multi-Mutation Library Generation and Screening

Plasmid DNAs from the screened single-mutation libraries were isolated using a Maxi-prep kit (Zymo Research) to generate multi-mutation libraries via SSM as described above, and using the DNA shuffling methodology as described previously^.24^ Input DNA from the VH and VL genes were amplified from the library plasmids using Kapa Hifi HotStart ReadyMix (Roche)^26^, and amplicons were randomly fragmented by introducing double stranded breaks with DNase I in the presence of MnCl_2_ at 15°C for 35-45 seconds. The resulting DNA fragments were reassembled using Platinum^TM^ Taq DNA polymerase (family A polymerase, Invitrogen) and AccuPrime^TM^ Pfx DNA polymerase (family B polymerase, Invitrogen), followed by re-amplification with Kapa Hifi HotStart ReadyMix (Roche). The separate shuffled-VH and shuffled-VL genes were combined into a single shuffled-VH:VL library via step-wise cloning^16^. Subsequently, yeast libraries were generated by homologous recombination with VL-BDP-VH expression cassette and pCT backbone as described previously^16,20,21^; library sizes exceeding 1E+07 were maintained for all multi-mutation libraries. 3E+07 yeast cells were screened in the first round of screening for all multi-mutation libraries.

Libraries from the first round of DNA shuffle mutagenesis were labeled with 1.25 nM biotinylated Peptide 22 (NANPNVDPNANPNVD) conjugated with streptavidin-PE (S21388, Invitrogen), and sorted for three rounds by FACS, as described above. Plasmids of the enriched populations after Round 3 were isolated, and SSM or DNA shuffling were performed to the VH and VL genes to add additional mutational diversity for the second round of DNA shuffle mutagenesis. Subsequently, yeast libraries were generated as described above. Libraries from second shuffle mutagenesis were labeled with monovalent antigen molecules, instead of antigen molecules pre-conjugated with streptavidin-PE (S21388, Invitrogen). Libraries were enriched against 0.5 nM of biotinylated Peptide 22 in the first round of FACS, 0.25 nM of biotinylated Peptide 22 in the second round of FACS, and 0.06 nM of biotinylated truncated-PfCSP (N-terminal domain stabilized truncated-PfCSP, NTDS_5/3)^7^ in the third round of FACS. NTDS_5/3 is a N-terminal domain stabilized truncated CSP, 3 with adjacent NPNV motifs. It was created by introducing four N-terminal domain stabilizing (NTDS) mutations, C25S and _66_KKNSR_70_ to _66_SSNSA_70_, to the 3D7 clone PfCSP, and reducing the extensive central repeat region by displaying five major repeat peptides (NANP) instead of 38, and three minor repeat peptides (NVDP) instead of four, while keeping the C-terminus unaltered^7,16^.

Because L9 is already a very high affinity molecule, depletion of antigen from solution can be observed at very low staining concentrations, for example during the later rounds of screening (**Figures S2-S3**). Antigen depletion effects limit the overall MFI observed in enriched populations for high-affinity antibodies like L9 at low antigen concentrations; still, affinity-gate sorting was used successfully to improve antibody affinity as we showed in this study.

### NGS and Bioinformatic Analysis of Sorted Yeast Display Libraries

We sequenced pre-sort libraries with Illumina MiSeq, which provided high sequence coverage. Due to the shorter length of the MiSeq reads, heavy and light antibody chains were sequenced separately. For sorted library populations with reduced diversity, the long-read Pacific Biosciences SMRT HiFi system was also used to sequence paired heavy and light antibody chains together. These two sets of sequencing data were first processed separately, and their information was mined together to identify enriched variants for experimental validation.

In the MiSeq data pipeline, paired end reads were first merged to assemble the entire heavy or light antibody chain via FLASH (v1.2.11).^30^ Then, the linked reads were quality filtered with FASTX-Toolkit (v0.0.14) to retain sequences where at least 90% of the bases had a Phred Quality Score of at least 30 (corresponding to 99.9% base call accuracy). IgBLAST (v1.16.0)^31^ was then used to annotate the antibody V(D)J genes and identify CDR3 sequences, with only productive sequence alignments retained in the output files.

For the Pacific Biosciences data, the raw sequence data was demultiplexed using Lima (v2.7.1). Once individual sample sequence files were recovered, the long reads were split into two reads, one with the heavy chain and the other with the light chain; pairing information was preserved via sequence headers. For separate heavy and light chain sequences, IgBlast (v1.16.0)^31^ was used to map V-(D)-J genes and identify CDR3 regions. Subsequently, paired heavy and light chain alignments were organized based on unique header information. Each heavy and light chain sequence was compared to the wild-type antibody sequence to identify substitution mutations. Enrichment analysis was performed using both MiSeq and Pacific Biosciences sequence data to identify antibody variants with improved antigen binding. The enrichment ratio (ER), calculated as variant frequency in a sorted library divided by the same variant frequency in the presort library, quantifies how much a variant has enriched after sorting for binding to an antigen^16,20,21,32^. Because the presort library was sequenced as separate heavy and light chain sequences, and so a Pacific Biosciences variant’s presort frequency was estimated as the heavy chain sequence’s MiSeq frequency multiplied by the light chain’s MiSeq frequency. ER was also calculated independently for the separate heavy and light chain variants. Heavy chain variants and light chain variants of interest were identified based on either high fractional representation or high ER, and Pacific Biosciences sorted library data were filtered for sequences with exact matches to candidates’ amino acid sequences. Nucleotide consensus sequences were generated from the filtered library data via USEARCH (v 5.2.236)^33^, enforcing a 90% sequence identity for clusters. Finally, full candidate antibody sequences were assembled by pairing the heavy and light chain consensus sequences together. In these Pacific Biosciences data, paired VH:VL sequences from FACS R3 with prevalence >0.1% and ER > 1 were classified as high-affinity variants.

### Antibody Expression and Purification

Antibody variable heavy chain and light chain sequences were codon optimized, synthesized and cloned into a VRC8400 (CMV/R expression vector)-based IgG1 vector as previously described (Kong et al, 2019).The variants were expressed by transient transfection in Expi293 cells (ThermoFisher Scientific, Waltham, MA) using Turbo293 transfection reagent (SPEED BioSystems, Gaithersburg, MD) according to the manufacturer’s recommendation. 50 μg plasmid encoding heavy-chain and 50 μg plasmid encoding light-chain variant genes were mixed with the transfection reagents, added to 100 ml of cells at 2.5 × 10^6^/ml, and incubated in a shaker incubator at 120 rpm, 37°C, 9% CO2. At 5 days post-transfection, cell culture supernatant was harvested and purified with a Protein A (GE Healthcare, Chicago, IL) column. The antibody was eluted using IgG Elution Buffer (ThermoFisher Scientific, Waltham, MA) and were brought to neutral pH with 1 M Tris-HCl, pH 8.0. Eluted antibodies were dialyzed against PBS overnight before use.

### Affinity Measurement of L9-variants by Biolayer Interferometry (BLI)

Antibody binding affinity to various ligands were measured using biolayer interferometry on an Octet Red384 instrument (fortéBio, Menlo Park, CA) with streptavidin capture biosensors (fortéBio) in solid black tilt-well 96-well plates (Greiner Bio-One, Kremsmünster, Austria). Assays were performed with agitation at 30°C. Immobilization of biotinylated fl_PfCSP_3D7, NTDS_5/3_CSP, fl_PfCSP_Kenya2-3 (GenBank accession codes AF540462.1 & AF540463.1)^34^, Pep21, Pep22, and Pep29 was performed for 5s, followed by a 60s baseline in buffer (PBS + 1% BSA). Association with IgG (serially diluted from 1000 to 1.3 nM) was done for 240s, followed by a dissociation step in buffer for 1200s. In all Octet measurements, parallel correction to subtract systematic baseline drift was carried out by subtracting the measurements recorded for a loaded sensor incubated in PBS. Data analysis was carried out using Octet software, version 9.0. Experimental data were fitted globally with a 1:1 Langmuir model of binding for all the antigens.

### Evaluation of Protective Potency of L9-variants Against Sporozoite Challenge in Mouse Model

Female 6- to 8-weeks old C57Bl/6J mice were obtained from The Jackson Laboratory (Bar Harbor, ME). All animals were maintained and cared for in accordance with the American Association for Accreditation of Laboratory Animal Care Standards. All mouse procedures were performed according to protocols approved by the Institutional Animal Care and Use Ethics Committees at Johns Hopkins University, Protocol number MO24H329.

To generate sporozoites, transgenic *P. berghei* (strain ANKA 676m1C11, MRA-868) expressing full-length *P. falciparum* CSP and a green fluorescent protein/luciferase fusion protein (Pb-PfCSP-GFP/Luc-SPZ or Pb-PfCSP-SPZ) were obtained from salivary glands of infected mosquitoes, as previously described^27^. Briefly, *Anopheles stephensi* mosquitoes were obtained and reared from a colony maintained at the insectary at Johns Hopkins University, Malaria Research Institute. Female mosquitoes were allowed to feed on 6- to 8-week-old female Swiss Webster mice infected with blood-stage Pb-PfCSP-GFP/LUC parasites. After infection, mosquitoes were maintained in an incubator at 19-20°C and supplied with a sterile cotton pad soaked in 10% sucrose, changed every 48 hrs. 20-23 days following mosquito infections, salivary glands were dissected and ground in 2% FBS-HBSS, SPZs were counted in a Neubauer chamber.

To assess the protective efficacy of anti-PfCSP mAbs *in vivo*, mAbs were diluted in sterile filtered 1x PBS and administered into the tail veins of female C57Bl/6J mice (The Jackson Laboratory, Bar Harbor, ME). After 2 hours, were intravenously challenged in the tail vein with 2,000 freshly harvested Pb-PfCSPGFP / Luc-SPZ in 2% FBS-HBSS (Thermo Fisher Scientific, Waltham, MA, USA).

To assess *Plasmodium* infection, mice were injected intraperitoneally (i.p.) with 100 μL d-luciferin (30 mg/mL, PerkinElmer, Waltham, MA), anesthetized with isoflurane and imaged with an IVIS® Spectrum *in vivo* imaging system (PerkinElmer) 5 minutes after luciferin injection. Parasite liver load was assessed 42 hours after challenge, whereas parasitemia was measured 6 days following challenge. Parasite load was quantified by analyzing a region of interest (ROI) in the upper abdominal region for liver stage, or whole animal for parasitemia; bioluminescence or the total flux (photons/second; p/s) was measured using the manufacturer’s software (Living Image 4.5, PerkinElmer).

### Cryo-EM Sample Preparation, Grid Preparation, and Data Collection

The sample for cryo-EM was prepared by mixing antigen binding fragment (Fab) to PfCSP (3D7) or PfCSP (Kenya2-3) at 2:1 molar ratio. The complex was incubated overnight at 4°C and flash frozen until ready for use. To prepare grids of the complex, 3 µL of protein sample was applied to freshly glow-discharged (easiGLow) C-flat grids (Protochips, CF1.2/1.3-3Au). Blotting was done using a Vitrobot Mark IV (Thermo-Fisher), with 5 s blotting time and 8 pN blotting force at 6 °C in 100% humidity. Grids were vitrified by plunging into liquid ethane and stored in liquid nitrogen before examination by cryo-EM. Images were recorded on a Glacios TEM (Thermo Fisher) at 200 kV and recorded at 36,000X magnification with a defocus range of −0.3 to −2.2 µm on K3 direct electron detector (Gatan) in super-resolution mode.

### Cryo-EM Data Processing and Refinement

Motion correction, contrast transfer function (CTF) estimation, particle picking, extraction, 2D classification, ab initio model generation, 3D refinements and local resolution estimation were carried out in cryoSPARC 3.3.1 (Punjani et al, 2017). The 3D reconstructions were performed using C1 symmetry for both the classes. The coordinates of L9-PfCSP structure, PDB entry (8EK1) was employed as initial model for fitting the sharpened cryo-EM map of the L9-PfCSP structures. Manual and automated model building were iteratively performed using Coot (Emsley & Cowtan, 2004) and real space refinement in Phenix (Adams et al, 2010) to accurately fit the coordinates to the electron density map. Molprobity (Davis et al, 2004) and EMRinger (Barad et al, 2015) was used to validate geometry and check structure quality. UCSF ChimeraX (Goddard et al, 2018) was used for map-fitting cross correlation calculation (Fit-in-Map tool) and for figure preparation.

### Quantification and Statistical Analysis

L9 variants were compared to an equivalent dose of L9_WT as a benchmark control, using a two-tailed ordinary one-way ANOVA test with Dunnett’s multiple test correction. Statistical significance between L9_yd19 and MAM01 was analyzed using a two-tailed unpaired t test. Statistical differences were estimated on log_10_(liver burden) values. To enable comparison of the correlation between parasite infection and antigen/epitope affinity across different studies, the liver burden of each group was normalized based on the median of the liver burden values from the untreated (max burden) mice in the same experiment. Pearson’s correlation coefficient (R) was calculated using the median normalized liver burden values with corresponding two-tailed P values. A linear regression was fit based on a Pearson’s correlation analysis. Data were plotted and graphed using GraphPad Prism, unless otherwise stated. P values ≤ 0.05 were considered significant as indicated in the figures.

## SUPPLEMENTARY FIGURE LEGENDS

**Figure S1.**
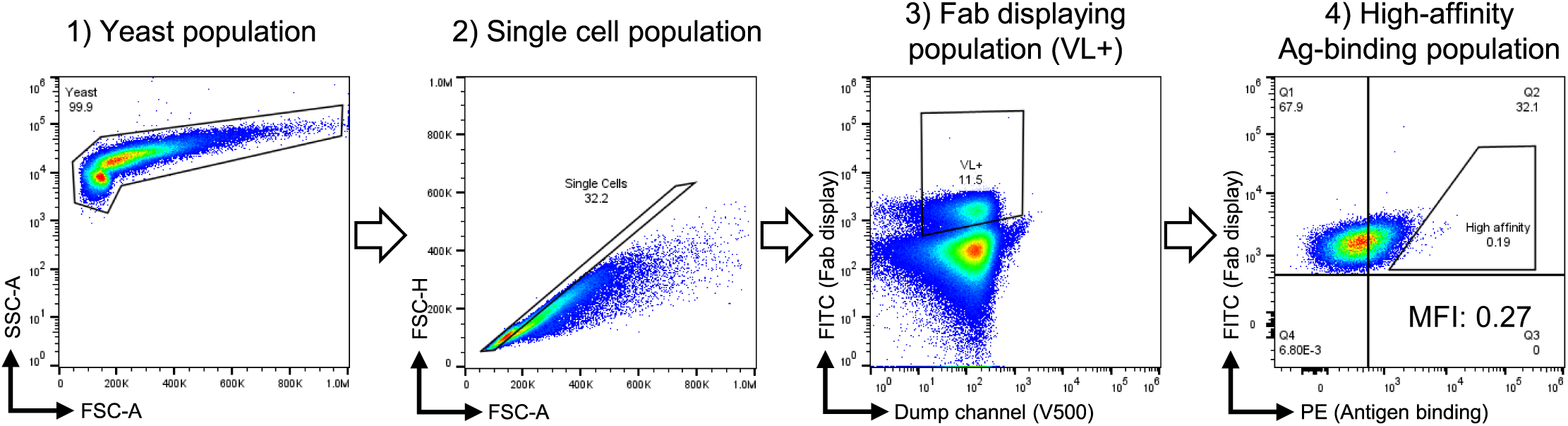
Example of flow cytometry gating for library sorts, related to Fig. 2. Overview of yeast display library gating strategy in the following order: 1) yeast population, 2) single cell population, 3) Fab-displaying population (VL+), and 4) high-affinity antigen-binding population. The mean fluorescence intensity ratio (MFI) is calculated as (Geomean _PE_ / Geomean _FITC_) to quantitatively represent the normalized affinity of Fab molecules.

**Figure S2.**
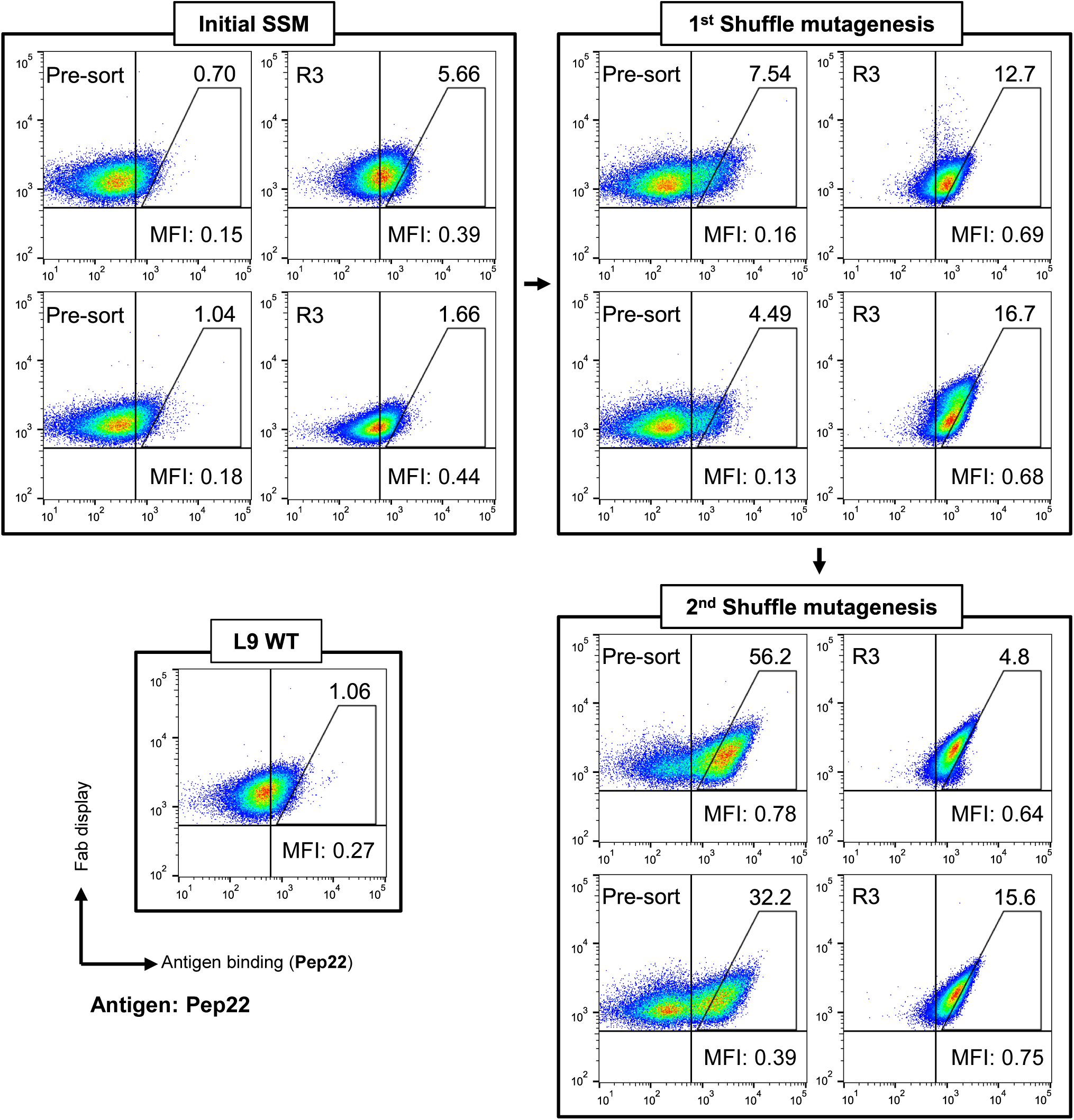
Selected flow cytometry plots labeled with Pep22, related to Fig. 2. Representative flow cytometry plots for pre-sort and FACS enriched libraries from across screening rounds labeled with 62.5 pM Pep22. An analytical high-affinity gate is drawn for comparison across libraries. The percentage of the high affinity population is shown in the upper-right quadrant. The mean fluorescence intensity ratio (MFI) is calculated as (Geomean _PE_ / Geomean _FITC_) to quantitatively represent the normalized affinity of Fab molecules.

**Figure S3.**
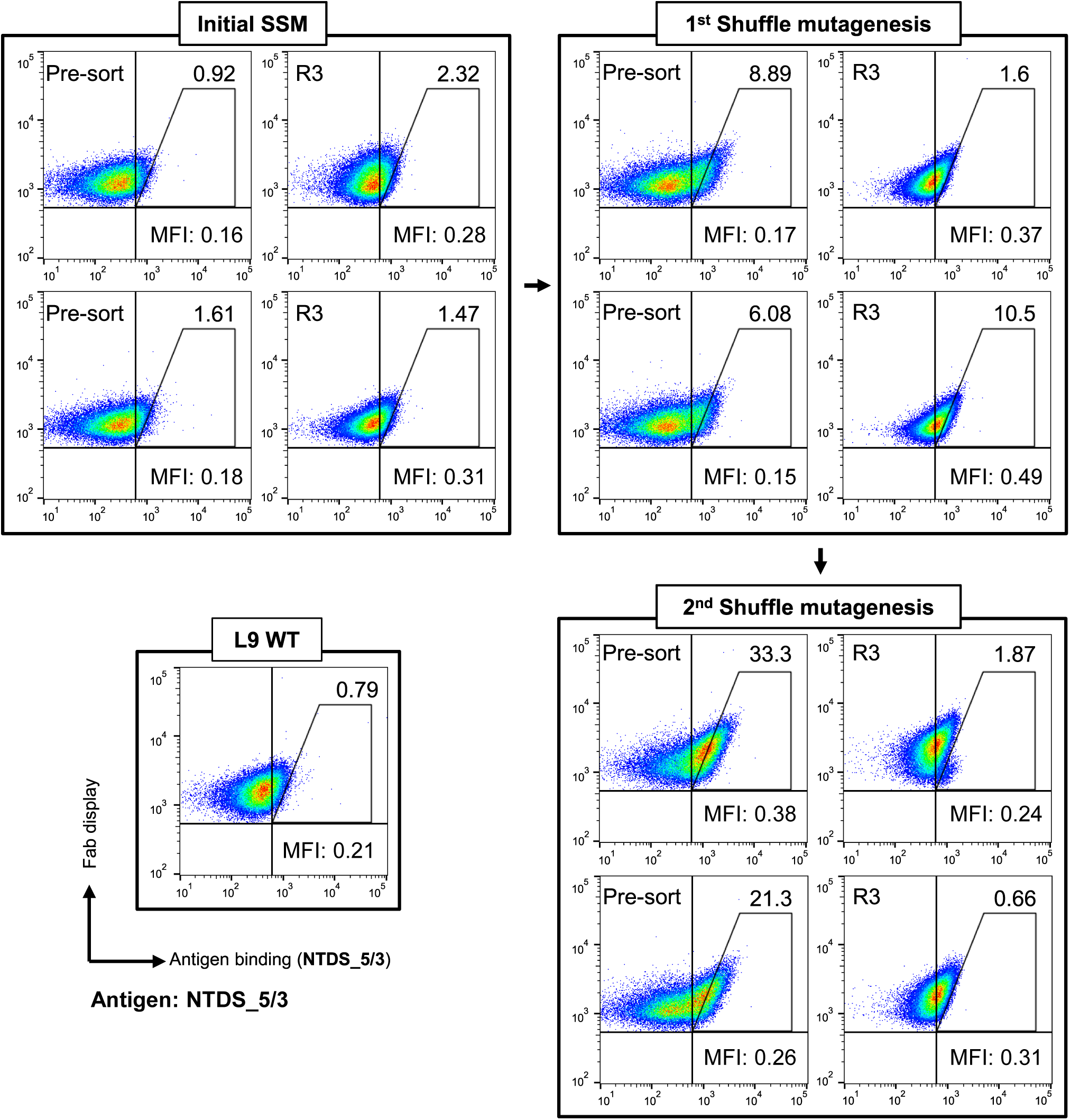
Selected flow cytometry plots labeled with NTDS_5/3, related to Fig. 2. Representative flow cytometry plots for pre-sort and FACS enriched libraries across screening rounds, labeled with 15.6 pM NTDS_5/3. An analytical high-affinity gate is drawn for comparison across libraries. The percentage of the high affinity population is shown in the upper-right quadrant. The mean fluorescence intensity ratio (MFI) is calculated as (Geomean _PE_ / Geomean _FITC_) to quantitatively represent the normalized affinity of Fab molecules.

**Figure S4.**
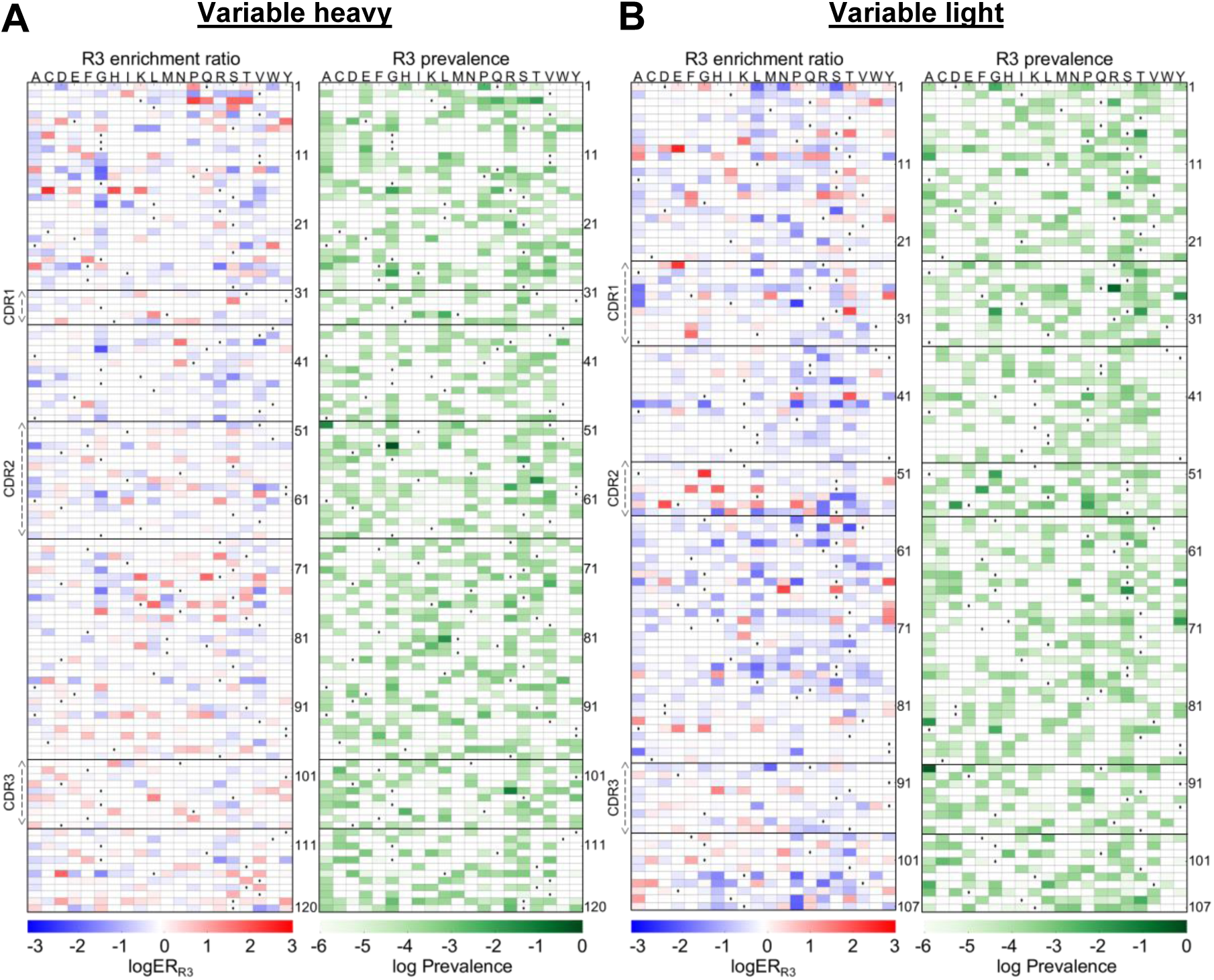
Bioinformatic analysis of paired VH:VL sequences from Round 3 (R3) of the 1^st^ shuffle mutagenesis, related to Fig. 3. The Round 3 (R3) enrichment ratio (left) and prevalence (right) for amino acid mutations identified from **(A)** VH sequences and **(A)** VH sequences and **(B)** VL sequences. Wild-type amino acids are indicated with dots (·). The amino acid residues followed template numbering.

**Figure S5.**
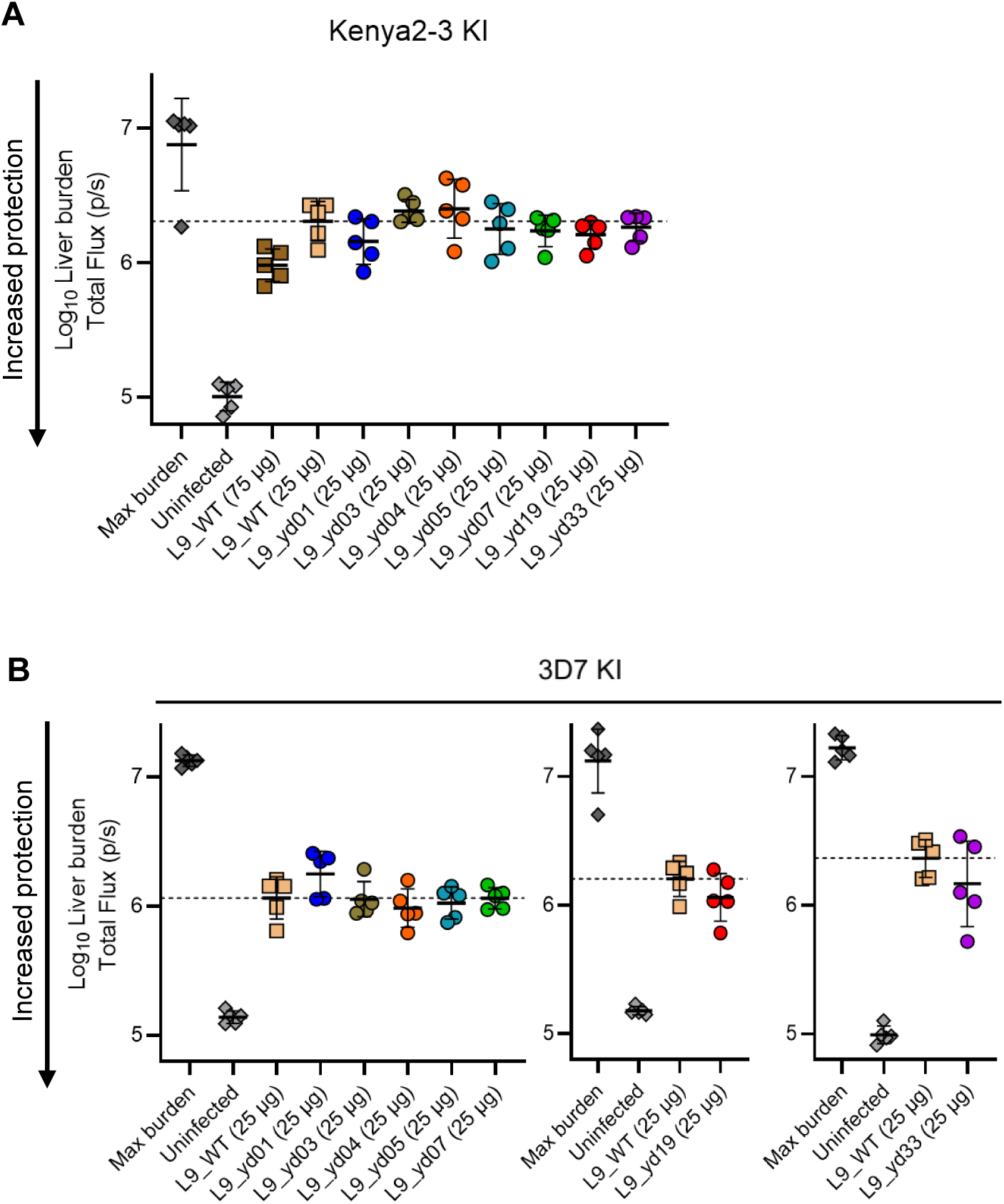
*In vivo* characterization of L9 variants in a mouse sporozoite challenge model, related to Fig. 4. Antibody variants were evaluated for protection against transgenic *P. berghei* sporozoites (PbSPZ) *in vivo* carrying either **(A)** Kenya2-3 sequence or **(B)** the 3D7 sequence. Each group consisted of five mice, with animals receiving the indicated antibody dose. Liver burden was quantified, with mean values ± standard deviation shown as bars. Statistical significance between L9_WT and each L9 variant group at 25 µg were determined using the ordinary one-way ANOVA test with a two-tailed P value calculation and a Dunnett’s multiple test correction. No L9 variants were statistically significant from L9_WT.

**Figure S6.**
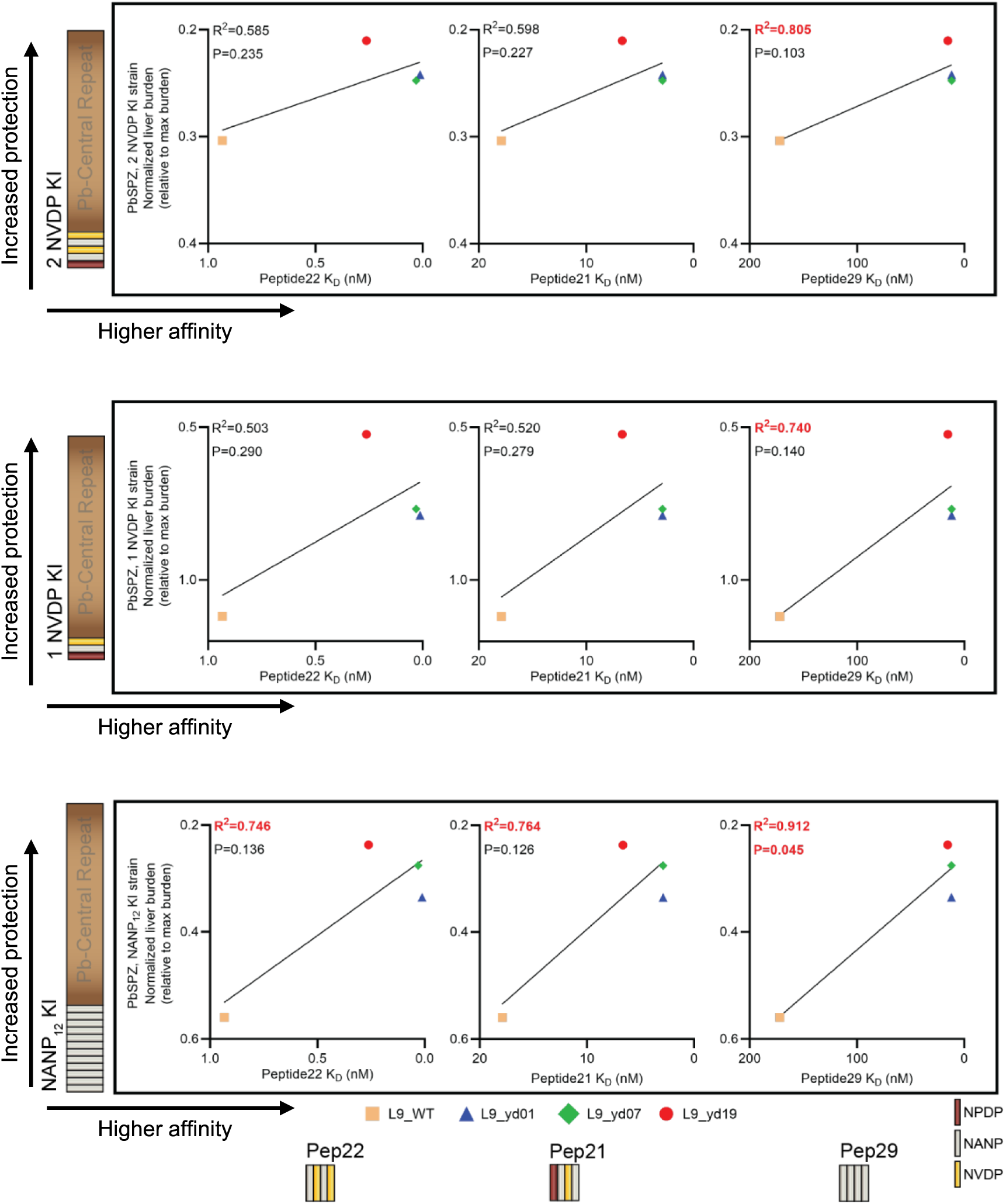
Correlation between the epitope-specific affinity and *in vivo* potency against transgenic KI strains, related to Fig. 5. Correlation between binding affinity to various peptide antigens (x-axis, linear scale) and normalized liver burden against various transgenic PbSPZs (y-axis, linear scale). Pearson’s correlation coefficient (R) was calculated using the median normalized liver burden values, with corresponding two-tailed P-values and 95% confidence intervals. A linear regression was fit based on a Pearson’s correlation analysis.

**Figure S7.**
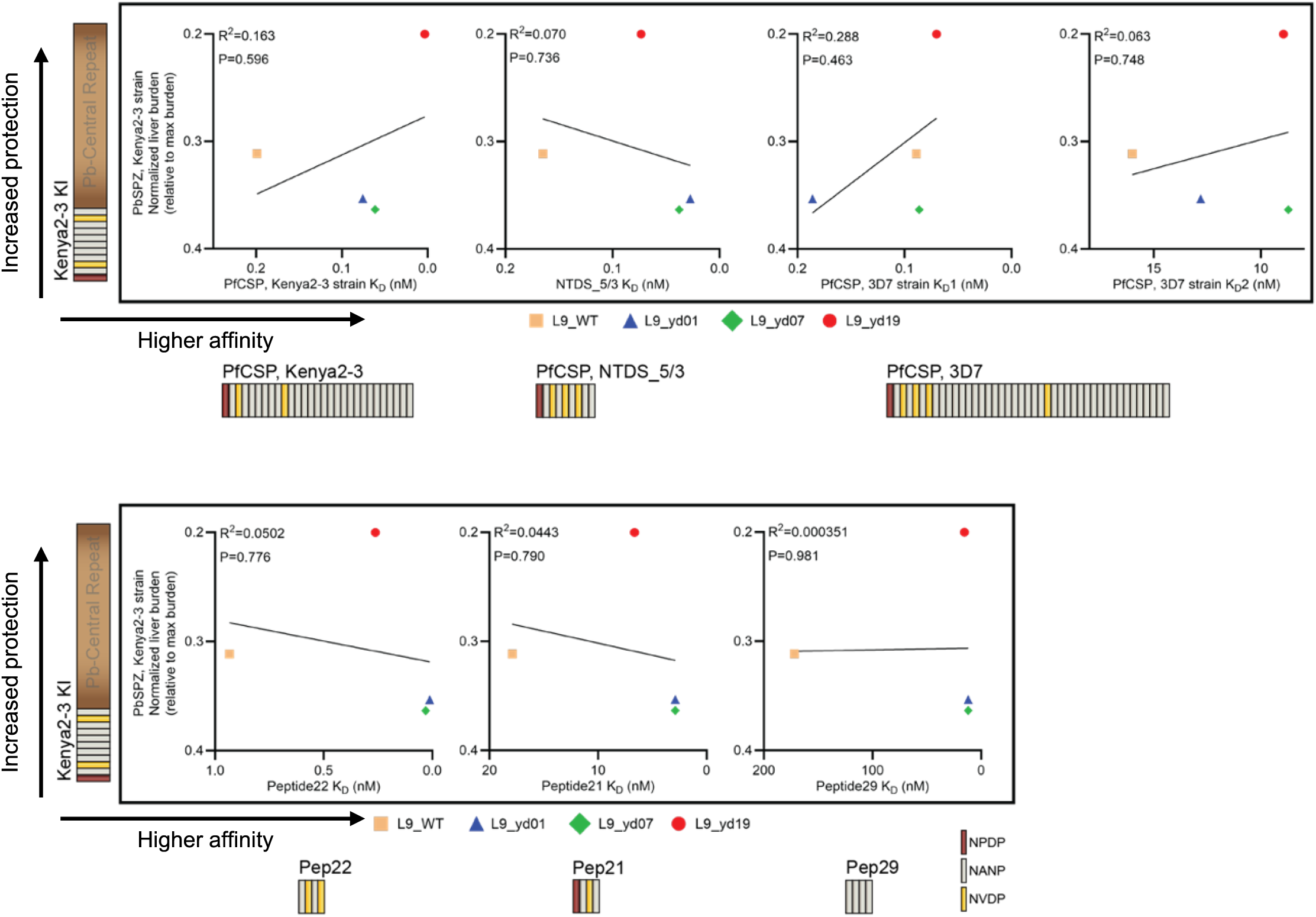
Correlation between the epitope-specific affinity and *in vivo* potency against the Kenya2-3 KI strain, related to Fig. 5. Correlation between binding affinity to various peptide antigens (x-axis, linear scale) and normalized liver burden against Kenya2-3 KI transgenic PbSPZ (y-axis, linear scale). Pearson’s correlation coefficient (R) was calculated using the median normalized liver burden values, with corresponding two-tailed P-values and 95% confidence intervals. A linear regression was fit based on a Pearson’s correlation analysis.

**Figure S8.**
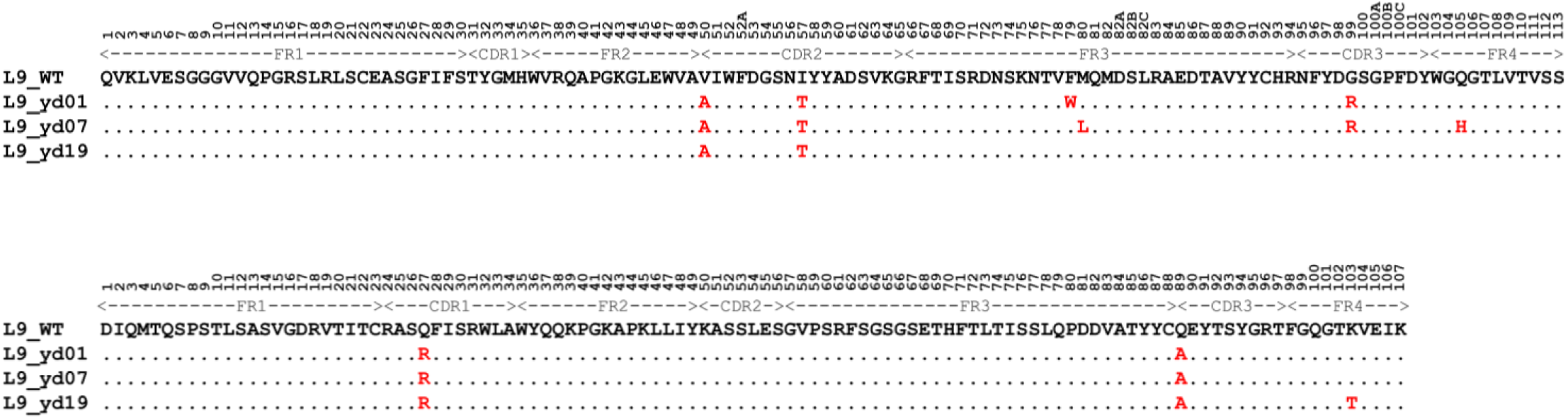
Sequence alignment of L9_WT and L9 variants, related to Fig. 6. Sequence alignment of L9_WT, L9_yd01, L9_yd07, L9_yd19, for the variable regions of the heavy chain (*upper*) and light chain (*lower*). Amino acid mutations identified in each L9-variant are indicated in red. Kabat numbering was used for the position of mutations.

## SUPPLEMENTARY TABLE LEGENDS

**Table S1.** Antibody variant mutations and affinity data, related to Fig. 4.

| mAb ID | VH mutations (template) | VH mutations (Kabat) | VL mutations (template) | VL mutations (Kabat) | Peptide22 K<sub>D</sub> (nM) | 3D7 fICSP K<sub>D</sub>1 (nM) | 3D7 fICSP K<sub>D</sub>2 (nM) |
| --- | --- | --- | --- | --- | --- | --- | --- |
| L9_WT |  |  |  |  | 0.934 | 0.089 | 16.00 |
| L9_yd01 | V50A \| I58T \| F80W \| G103R | V50A \| I57T \| F79W \| G99R | Q27R \| Q89A | Q27R \| Q89A | 0.012 | 0.186 | 12.82 |
| L9_yd02 | V50A \| I58T \| M81L \| G103R \| G113D | V50A \| I57T \| M80L \| G99R \| G106D | Q27R \| Q89A | Q27R \| Q89A | 1.071 | 1.025 | 692.80 |
| L9_yd03 | V50A \| I58R \| M81L \| G103R | V50A \| I57R \| M80L \| G99R | Q27R \| Q89A | Q27R \| Q89A | 0.068 | 0.126 | 10.67 |
| L9_yd04 | R16G \| I28G \| V50A \| G55S \| I58T \| G103R | R16G \| I28G \| V50A \| G54S \| I57T \| G99R | Q27R \| Q89A | Q27R \| Q89A | 0.017 | 0.218 | 11.42 |
| L9_yd05 | V50A \| G55D \| I58T \| M81L \| G103R | V50A \| G54D \| I57T \| M80L \| G99R | Q27R \| Q89A | Q27R \| Q89A | 0.036 | 0.166 | 11.32 |
| L9_yd06 | V50A \| I58T \| M81L \| R98G \| G103R | V50A \| I57T \| M80L \| R94G \| G99R | Q27R \| Q89A | Q27R \| Q89A | 20.110 | 0.623 | 43.98 |
| L9_yd07 | V50A \| I58T \| M81L \| G103R \| Q112H | V50A \| I57T \| M80L \| G99R \| Q105H | Q27R \| Q89A | Q27R \| Q89A | 0.030 | 0.086 | 8.71 |
| L9_yd08 | A49T \| V50A \| I58T \| M81L \| G103R | A49T \| V50A \| I57T \| M80L \| G99R | Q27R \| Q89A | Q27R \| Q89A | 0.022 | 0.089 | 9.21 |
| L9_yd09 | A24T \| V50A \| I58T \| M81L \| G103R | A24T \| V50A \| I57T \| M80L \| G99R | Q27R \| Q89A | Q27R \| Q89A | 0.015 | 0.129 | 9.48 |
| L9_yd10 | S7F \| V50A \| I58T \| M81L \| G103R | S7F \| V50A \| I57T \| M80L \| G99R | Q27R \| Q89A | Q27R \| Q89A | 0.034 | 0.158 | 9.28 |
| L9_yd11 | V50A \| I58T \| F80V \| G103R | V50A \| I57T \| F79V \| G99R | Q27R \| Q89A | Q27R \| Q89A | 0.134 | 0.106 | 7.83 |
| L9_yd12 | A49V \| V50A \| I58T \| G103R | A49V \| V50A \| I57T \| G99R | Q27R \| Q89A | Q27R \| Q89A | 0.121 | 0.194 | 19.65 |
| L9_yd13 | M34V \| V50A \| I58T \| G103R | M34V \| V50A \| I57T \| G99R | Q27R \| Q89A | Q27R \| Q89A | 0.037 | N/A | 1.30 |
| L9_yd14 | I28G \| G103R | I28G \| G99R | Q27R \| Q89A | Q27R \| Q89A | 0.272 | 0.210 | 17.17 |
| L9_yd15 | I28R \| M81L \| G103R | I28R \| M80L \| G99R | Q27R \| Q89A | Q27R \| Q89A | 0.224 | 0.199 | 19.47 |
| L9_yd16 | S21T \| F53G \| Y59R \| F80V \| G103R | S21T \| F52AG \| Y58R \| F79V \| G99R | Q27R \| Q89A | Q27R \| Q89A | 0.110 | 0.081 | 12.94 |
| L9_yd17 | I28R \| A40T \| M81L \| G103R | I28R \| A40T \| M80L \| G99R | Q27R \| Q89A | Q27R \| Q89A | 0.154 | 0.235 | 21.79 |
| L9_yd18 | V50A \| G55S \| I58T \| G103R | V50A \| G54S \| I57T \| G99R | Q27R \| S56R \| Q89A | Q27R \| S56R \| Q89A | 0.033 | 0.252 | 13.30 |
| L9_yd19 | V50A \| I58T | V50A \| I57T | Q27R \| Q89A \| K103T | Q27R \| Q89A \| K103T | 0.262 | 0.070 | 8.94 |
| L9_yd20 | R16G \| F53G \| I58T \| G103R | R16G \| F52AG \| I57T \| G99R | T10L \| Q27R \| Q89A \| V104L | T10L \| Q27R \| Q89A \| V104L | 0.264 | 0.404 | 32.48 |
| L9_yd21 | F53G \| G103R | F52AG \| G99R | Q27R \| A51T \| Q89A | Q27R \| A51T \| Q89A | 0.238 | 0.461 | 33.93 |
| L9_yd22 | V50A \| I58T \| G103R | V50A \| I57T \| G99R | A25P \| Q27R \| Q89A | A25P \| Q27R \| Q89A | 0.157 | 0.307 | 18.06 |
| L9_yd23 | I28G \| V50A \| I58T \| G103R | I28G \| V50A \| I57T \| G99R | P8S \| V19L \| Q27R \| L54R \| Q89A | P8S \| V19L \| Q27R \| L54R \| Q89A | 0.077 | 0.287 | 17.44 |
| L9_yd24 | S7W \| I28G \| M81L \| G103R | S7W \| I28G \| M80L \| G99R | Q27R \| K42Q \| Q89A | Q27R \| K42Q \| Q89A | 0.191 | 0.275 | 17.92 |
| L9_yd25 | V50A \| I58T \| G103R | V50A \| I57T \| G99R | S9L \| Q27R \| Q89A \| V104L | S9L \| Q27R \| Q89A \| V104L | 0.091 | 0.149 | 9.52 |
| L9_yd26 | I58T \| M81L \| G103R | I57T \| M80L \| G99R | Q27R \| Q89A | Q27R \| Q89A | 0.202 | 0.194 | 14.82 |
| L9_yd27 | V50A \| I58T \| G103R | V50A \| I57T \| G99R | I2T \| Q27R \| Q89A | I2T \| Q27R \| Q89A | 0.136 | 0.214 | 9.30 |
| L9_yd28 | V50A \| I58T \| D73R \| M81L \| G103R | V50A \| I57T \| D72R \| M80L \| G99R | Q27R \| Q89A | Q27R \| Q89A | 0.073 | N/A | 0.74 |
| L9_yd29 | V50A \| I58T \| G103R | V50A \| I57T \| G99R | Q27R \| G57H \| Q89A | Q27R \| G57H \| Q89A | 0.044 | 0.132 | 11.22 |
| L9_yd30 | S25D \| V50A \| I58T \| G103R | S25D \| V50A \| I57T \| G99R | Q27R \| L54R \| Q89A | Q27R \| L54R \| Q89A | 0.054 | 0.354 | 15.84 |
| L9_yd31 | I28R \| Y59R \| T69E \| M81L \| G103R | I28R \| Y58R \| T68E \| M80L \| G99R | Q27R \| Q89A | Q27R \| Q89A | 0.148 | 0.219 | 20.38 |
| L9_yd32 | V50A \| I58T \| M81L \| G103R | V50A \| I57T \| M80L \| G99R | Q27R \| Q89A \| K103R \| V104W \| E105R \| I106S | Q27R \| Q89A \| K103R \| V104W \| E105R \| I106S | No expression yield | No expression yield | No expression yield |
| L9_yd33 | S25T \| V50A \| I58T \| G103R | S25T \| V50A \| I57T \| G99R | Q89A | Q89A | 0.064 | 0.239 | 14.82 |
| L9_yd34 | G26D \| V50A \| I58T \| M81L \| G103R | G26D \| V50A \| I57T \| M80L \| G99R | Q27R \| Q89A | Q27R \| Q89A | 0.099 | 0.317 | 16.28 |
| L9_yd35 | I28R \| A40T \| M81L \| V93M \| G103R | I28R \| A40T \| M80L \| V89M \| G99R | Q27R \| Q89A | Q27R \| Q89A | 0.194 | 0.218 | 20.47 |
| L9_yd36 | V50A \| I58T \| G103R | V50A \| I57T \| G99R | T10M \| Q89A | T10M \| Q89A | 0.118 | 0.151 | 9.98 |
| L9_yd37 | K43R \| F53G \| G103R | K43R \| F52AG \| G99R | S9P \| Q27R \| L33V \| Q89A | S9P \| Q27R \| L33V \| Q89A | 0.213 | 0.368 | 20.68 |

**Table S2.**
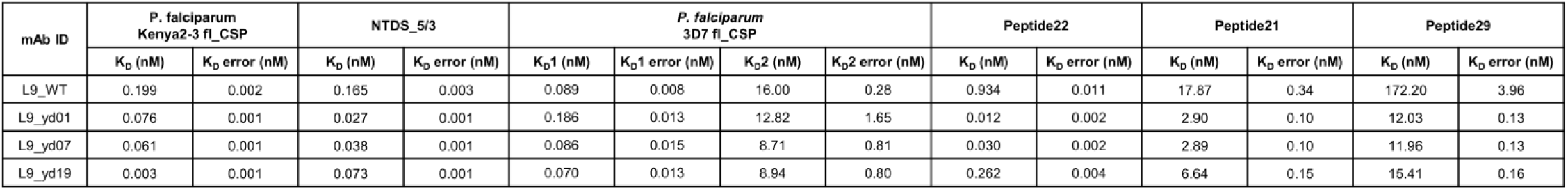
Affinity data of L9_WT and select L9 variants., related to Figs. 5, S6, and S7.

